# Spliceosomal mutations decouple 3′ splice site fidelity from cellular fitness

**DOI:** 10.1101/2023.01.12.523824

**Authors:** Kevin R. Roy, Jason Gabunilas, Dean Neutel, Michelle Ai, Joyce Samson, Guochang Lyu, Guillaume F. Chanfreau

**Author notes:** Current address: Department of Genetics, Stanford University School of Medicine, Stanford, California, United States of America (Kevin R. Roy). These authors contributed equally (Kevin R. Roy, Jason Gabunilas).

## Abstract

The fidelity of splice site selection is thought to be critical for proper gene expression and cellular fitness. In particular, proper recognition of 3′-splice site (3′SS) sequences by the spliceosome is a daunting task considering the low complexity of the 3′SS consensus sequence YAG. Here we show that inactivating the near-essential splicing factor Prp18p results in a global activation of alternative 3′SS, many of which harbor sequences that highly diverge from the YAG consensus, including some highly unusual non-AG 3′SS. We show that the role of Prp18p in 3′SS fidelity is promoted by physical interactions with the essential splicing factors Slu7p and Prp8p and synergized by the proofreading activity of the Prp22p helicase. Strikingly, structure-guided point mutations that disrupt Prp18p-Slu7p and Prp18p-Prp8p interactions mimic the loss of 3′SS fidelity without any impact on cellular growth, suggesting that accumulation of incorrectly spliced transcripts does not have a major deleterious effect on cellular viability. These results show that spliceosomes exhibit remarkably relaxed fidelity in the absence of Prp18p, and that new 3′SS sampling can be achieved genome-wide without a major negative impact on cellular fitness, a feature that could be used during evolution to explore new productive alternative splice sites.

## Introduction

Accurate splice site selection is critical for proper gene expression, as errors in splicing are known to cause a large number of genetic diseases and cancer^1^. However, proper splice site selection offers a major mechanistic challenge to the spliceosome in part because of the simplicity of splicing signals, as multiple near-consensus “incorrect” sites often exist in the vicinity of the “correct” site. This is especially problematic for 3′-splice sites (3′-SS) given its very simple consensus sequence YAG. Although the 3′-SS is typically selected prior to the first chemical step during metazoan splicing^2^, the second catalytic step of pre-mRNA splicing offers additional opportunity to further recognize the appropriate 3′-SS. Following an ATP-dependent rearrangement of the spliceosome after the first catalytic step^34^, Slu7p, Prp18p, and Prp22p sequentially join the spliceosome to facilitate the second catalytic step ^5,6^. Prp18p and Slu7p are necessary for the efficient docking of the 3′SS into the spliceosomal active site^7^, while Prp22p remodels pre-mRNA to promote sampling of alternative 3′SS^8^, and proofreads the ligated exons in an ATP-dependent manner following the second transesterification reaction^9^. Previous work using splicing reporters has shown that Prp18p plays a crucial role in stabilizing the interactions between exonic nucleotides and nucleotides of the U5 snRNA loop1^10^. Exonic nucleotides facilitate selection of the correct 3′SS especially when the function of Prp18p is partially compromised^11^. The role of Prp18p in stabilizing splicing intermediates for the second step of splicing ^10,11^ and its position near the 3′SS during the second step ^4,7^ suggests that it may also contribute to the fidelity of 3′SS selection. However, the global role of Prp18p in promoting splicing fidelity has not been investigated. To assess the impact of Prp18p on 3′SS selection genome-wide, we performed transcriptome analysis of cells lacking Prp18p and found that the loss of Prp18p resulted in a global loss of splicing fidelity and a widespread activation of 3′SS whose sequence deviate from the consensus. We show that the role of Prp18p in promoting 3′SS fidelity depends on its recruitment by Slu7p and involves positioning of Prp18p by the RNase H domain of Prp8p. We show that the role of Prp18p in promoting 3′-SS fidelity is synergized by the proofreading activity of the Prp22p helicase. Genetic analyses revealed that mutations of Prp18p and of its partners that affect 3′SS fidelity do not impact cellular fitness. These results show that a partial loss of splicing fidelity can be tolerated, possibly in order to explore novel splice sites and splice isoforms, revealing a possible mechanism to increase RNA isoforms diversity.

## Results

Prp18p was previously shown to suppress the usage of an unusual AUG 3′SS in the *GCR1* pre-mRNA^12^. However, the genome-wide role of Prp18p in promoting proper 3′SS selection is currently unknown. To globally analyze the impact of Prp18p on splicing fidelity, we performed RNA-seq on cells expressing wild-type spliceosomes or lacking Prp18p (*prp18*Δ) in a genetic background deficient for nonsense-mediated mRNA decay (NMD) due to the inactivation of the NMD helicase Upf1p (Fig.1a). The pseudo-wild-type *upf1*Δ genetic context was chosen to stabilize alternatively spliced mRNAs harboring premature termination codons (PTCs) ^12^, allowing a more direct analysis of the spliced transcript production without confounding effects of differential isoforms stability. We obtained 60-100 million uniquely mapped 100bp reads for each of the three replicates for each strain. 2-3% of all reads mapped with gapped alignments, and 93-95% of the gapped alignments corresponded to annotated splicing junctions.

**Figure 1.**
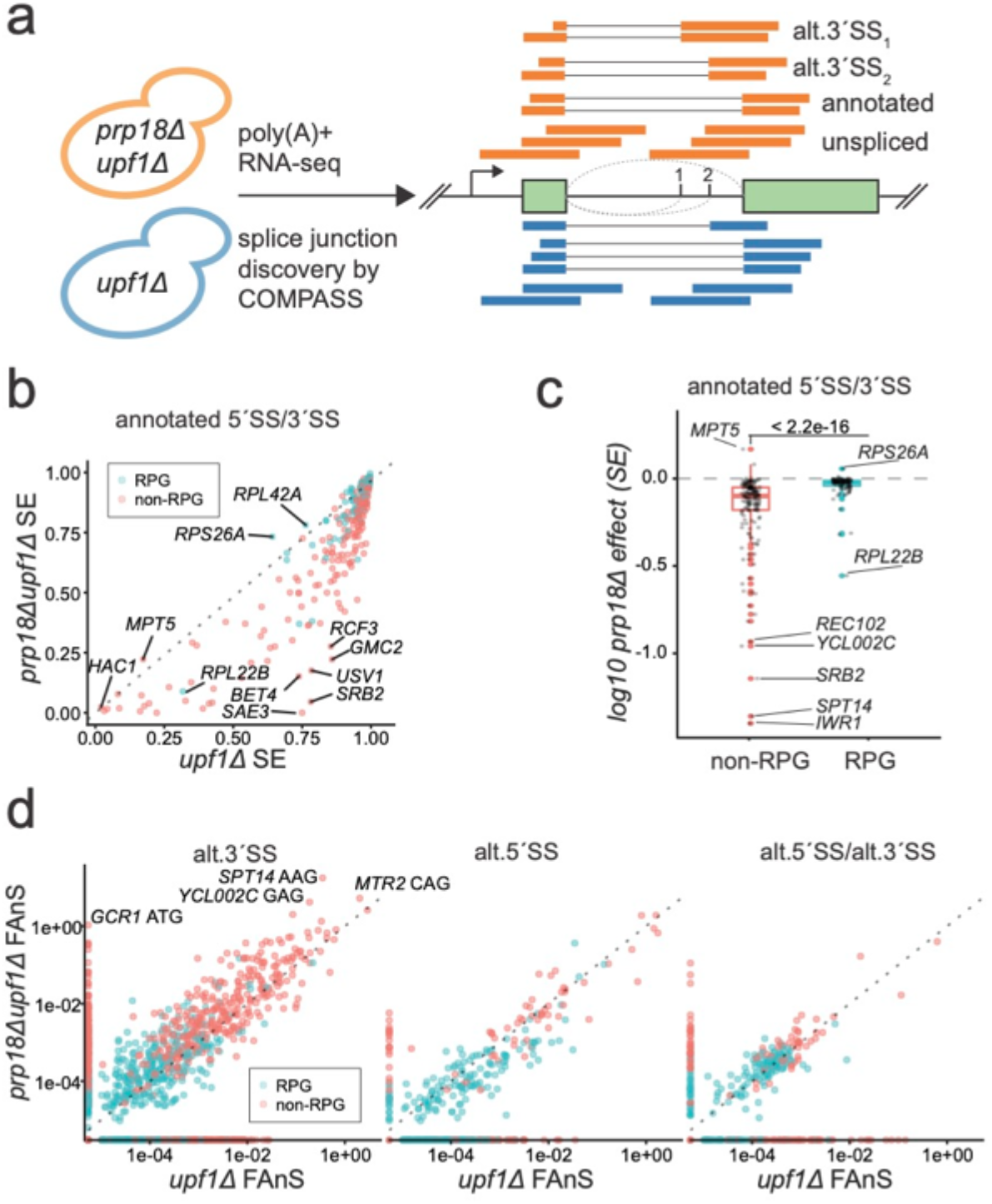
Global impact of Prp18p on annotated and alternative pre-mRNA splicing. (a) Diagram describing the workflow for identifying alternative splice junctions accumulating in prp18Δupf1Δ versus upf1Δ. Alignments generated by several annotation guided-aligner (eg. STAR) or annotation/motif agnostic aligners (eg BBMap) are processed by COMPASS to identify the optimal alignment for each read with a gapped alignment. (b) Splicing efficiency (SE) in prp18Δupf1Δ versus upf1Δ for annotated introns with ribosomal protein genes (RPGs) in blue and non-RPGs in red. (c) Effect of Prp18 inactivation of the splicing efficiency (SE) of intron containing genes separated in non-RPGs (red) and RPGs (blue) classes. Shown is a boxplot of the prp18Δ effect on SE (prp18Δupf1Δ SE/ upf1Δ SE) for annotated splice junctions. Log10 less than zero represents decreased splicing in the absence of Prp18p. (d) Fraction of annotated splicing (FAnS) for ann.5′SS/alt.3′SS (left), alt.5′SS/ ann.3′SS (middle), or alt.5′SS/alt.3′SS (right). Points on the axes indicate junctions for which splicing is not detected in the opposing strain (note the log scale). (d) Density distributions of total FAnS associated with each annotated intron for alt. 3′SS (left) and alt. 5′SS (right) with different density functions for RPG vs. non-RPG and prp18Δupf1Δ vs. upf1Δ. (e) The total number of alternative junctions observed as a function of minimum FAnS (with ≥5 reads across the entire dataset) and split into 3 lines according to alt. junction type. The horizontal line depicts the 1/1000 FAnS threshold.

We first analyzed the splicing efficiency (SE) for each annotated splice junction in the presence and absence of Prp18p (see Materials and Methods). Consistent with a previous analysis using tiling microarrays^13^, inactivation of Prp18p significantly reduced SE genome-wide (Fig.1b). As described in previous studies ^14,15^, ribosomal protein genes (RPGs) pre-mRNAs exhibited higher SE relative to non-RPGs in the wild-type strain at the annotated splice sites (Fig 1b). To further assess the impact of Prp18p on splicing, we calculated a *prp18*Δ-effect index for SE, where a positive *prp18*Δ effect indicates increased splicing in *prp18*Δ, and negative indicates decreased splicing (see Methods). As expected, the loss of Prp18 had an overall negative impact on SE for all annotated junctions (Fig.1b, 1c, Fig S1). However, there was a much greater negative impact on non-RPGs pre-mRNAs SE relative to RPGs (Fig.1c), showing that the splicing of RPG pre-mRNAs is more resilient to the loss of Prp18p than that of non-RPGs.

Next, we analyzed alternative splice junctions that arise in the absence of Prp18p. We noticed that STAR^16^, a commonly used RNA-seq aligner failed to map reads correctly for an alternative 3′SS that used non-canonical 3′SS sequences (e.g. AUG for the *MUD1* intron). By contrast, we found that BBMap [https://sourceforge.net/projects/bbmap/, https://www.osti.gov/biblio/1241166], a general-purpose aligner agnostic to splice motifs, was able to map these reads correctly but often failed to map reads with short overhangs at annotated junctions where STAR performed well. To combine the strengths of each aligner, we systematically compared STAR and BBMap alignments for each read harboring a potential splice-junction and selected the alignment exhibiting the fewest mismatches to the reference. We further removed likely false positive junctions through quality filtering of the STAR-BBMap integrated set of junctions to obtain a high-confidence set of alternative and novel junctions.

To analyze how splicing patterns at individual genes shift in the absence of Prp18p, we calculated the Fraction of Annotated Splicing (FAnS) for all alternative splicing junctions in the presence and absence of Prp18. The FAnS ratio reports the relative abundance of an alternative splicing event compared to the canonical, annotated main spliced isoform. RPGs pre-mRNAs constitute 90% of *S.cerevisiae* intron-containing mRNAs by transcript abundance^14^ even though they represent only 1/3 of all intron-containing genes (ICGs). Overall, RPGs exhibited *c.a*. 6-fold more annotated junctions reads than non-RPGs, but only 1.63-fold more alternative junctions reads in RPGs versus non-RPGs in the *upf1*Δ strain, and equal reads (1.01:1) in the *upf1*Δ*prp18*Δ strain (Fig S1). These observations show that the alternative splicing events detected do not simply arise due to spliceosomal noise that scales with expression levels, but are governed by specific features in ICGs that control splicing fidelity, as suggested previously^17^. Consistent with this, alternative junctions in non-RPGs showed significantly higher FAnS than those in RPGs (Fig 1d). Importantly, the FAnS calculated from our sequencing data set showed a high level of concordance with the FAnS calculated from a study examining alternative AG sites usage in a *upf1*Δ strain^17^ (Fig S2).

The absence of Prp18p globally activated alternative 3′SS usage and inhibited alternative 5′SS usage relative to annotated sites (Fig 1d). In particular, the absence of Prp18p revealed a large number of alternative 3′ SS for non-RPG transcripts which were virtually indetectable in the *upf1*Δ strain (Fig.1d, left panel), including the *GCR1* AUG 3′ SS site previously described (Kawashima et al., 2014). While for many splicing events, the increase in FAnS is largely driven by a decrease in annotated SE, many splice sites exhibited an increased SE in the absence of Prp18p (8 annotated sites, 294 alt.3′SS, 15 alt.5′SS, 47 alt.5′SS/alt.3′SS; Fig S3). Remarkably, 9 splice isoforms that used alternative 3′SS in 6 genes exhibited greater spliced mRNA abundance than the mRNA produced from the annotated sites (*GCR1*, *MTR2*, *REC102*, *REC107*, *SPT14*, *YCL002C)*, and 15% of all introns (42/280) exhibited at least 10% alt.3′SS usage. These data show that Prp18p promotes global recognition of *bona fide* 3′SS over competing alternative sites.

### Widespread activation of RAG, BG, and HAU 3′SS in the absence of Prp18p

To identify the genomic determinants for alternative 3′SS activation in the absence of Prp18p, we analyzed FAnS as a function of the sequence of these alternative 3′SS sequence. These 3′SS motifs could be broadly divided into YAG, RAG, BG (where B = C, G, or U), and non-G motifs., with roughly equal numbers of junctions (126 YAG, 166 RAG, 156 BG, 115 non-G, 6-reads minimum, Fig 2a,2b). The majority of these sites had not been detected in previous transcriptome-wide analyses of alternative splicing, likely a consequence of the low usage of these sites when Prp18p is functional and because of biases towards AG/AC introns in the mapping algorithms previously used^12,17–19^. Strikingly *prp18*Δ spliceosomes tended to specifically activate alternative 3′SS harboring non-YAG/non-AAG motifs (Fig 2a,2b). GAG and BG 3′SS were the most upregulated in the absence of Prp18p, with most alternative YAG sites instead being down-regulated (Fig 2a,2b). We confirmed the activation of the non-canonical 3′SS by performing targeted RT-PCR on selected transcripts, including non-canonical 3′SS UGG in *UBC12*, CUG in *MAF1*, AUG in *MUD1*, ACG in *PHO85* and UUG in *SPT14* (Fig 2c). We also confirmed usage of a highly unusual CAU 3′SS for *NYV1* (see below) For *SPT14* and *YCL002C*, RT-PCR confirmed the near-complete loss of the annotated spliced isoforms with concomitant activation of alternative upstream RAG sites (Fig 2c).

**Figure 2.**
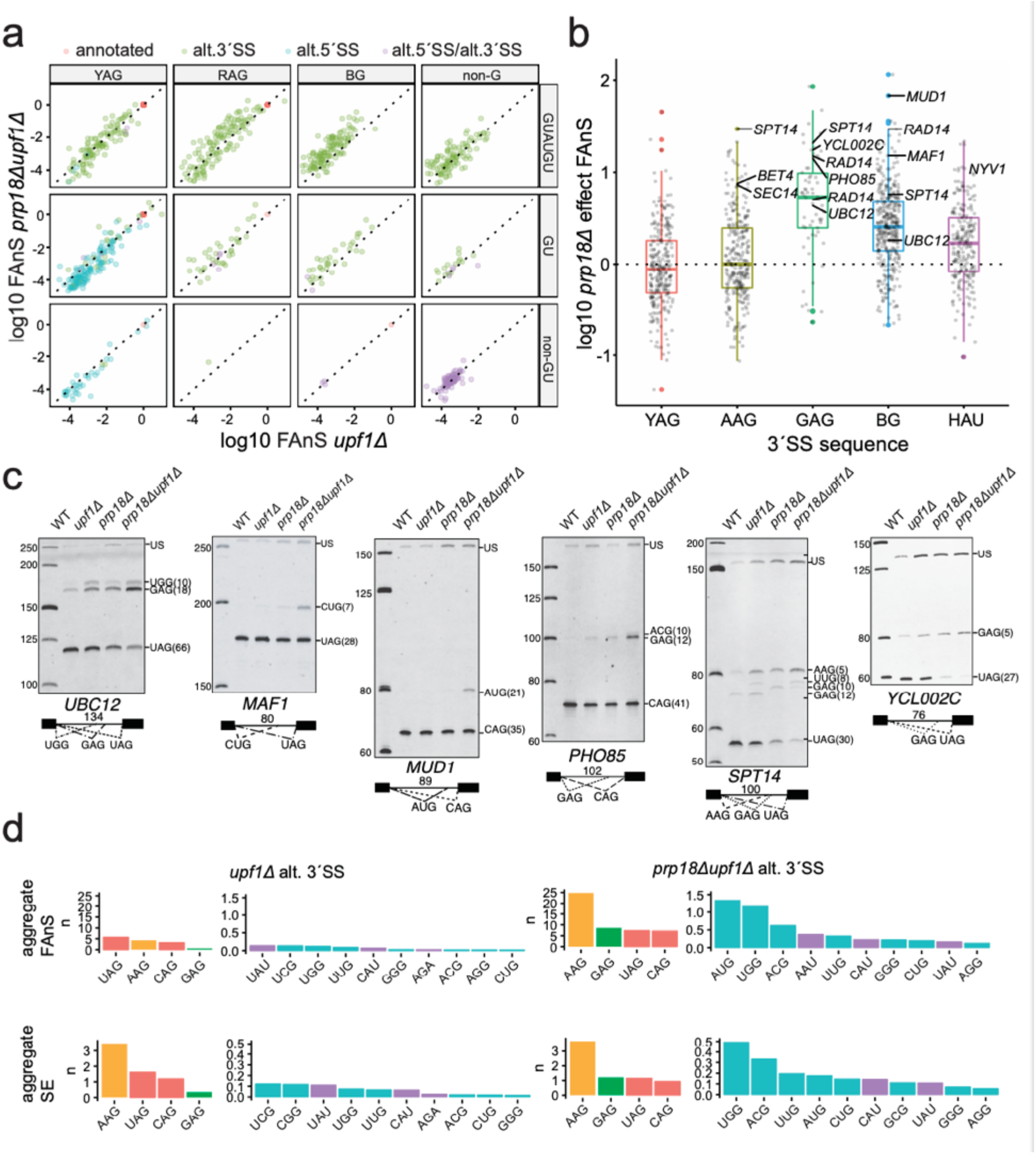
Widespread activation of non-canonical 3′SS upon inactivation of Prp18p. (a) Fraction of annotated splicing (FAnS) and (b) boxplots of the *prp18*Δ effect (*prp18*Δ*upf1*Δ FAnS/ *upf1*Δ FAnS) for different 3′SS motif class. (c) Selected junctions validated by RT-PCR with Cy3-labeled primers and polyacrylamide gel electrophoresis. The numbers in each intron diagram represent the length of the annotated intron. The numbers beside each 3′SS sequence on the gel represent distance (in bp) from the branchpoint (BP) to the indicated 3′SS. Note that each of these alternative junctions as well as those in Fig S8 is labeled in the *prp18*Δ effect boxplots in panel b. (d) Aggregate usage of each motif class calculated as the sum of FAnS or SE for all alternative sites in each class. Salmon = YAG; Green = GAG; Orange = AAG; Cyan = BG; Purple = HAU.

We next analyzed the aggregate usage for each 3′SS sequence genome-wide by summing SE and FAnS across transcripts, and found AAG to be the most utilized alternative 3′SS sequence by total SE (Fig 2d). Although the overall SE for AAG was roughly the same in both strains, the FAnS at AAG sites was substantially higher in the absence of Prp18p, suggesting a preferential loss of annotated 3′SS usage in the absence of Prp18p (Fig 2d, S4). While AAG spliced junctions outnumber GAG junctions, the SE for GAG increased by a larger extent in the absence of Prp18p relative to AAG (Fig 2d). AUG, ACG, UGG, and UUG were the most efficiently used non canonical BG sites, while CAU and UAU were the most efficiently used non-G sites (Fig 2d). In summary, these results demonstrate that in the absence of Prp18p, the spliceosome can utilize 3′SS lacking the consensus A or G nucleosides that are the hallmark of most 3′SS. In other words, spliceosomes exhibit remarkably relaxed fidelity in the absence of Prp18p, with a reduced ability to discriminate YAG from non-YAG sequences.

### Prp18p prevents activation of branchpoint-proximal 3′SS and non-canonical 3′SS

We next analyzed the features of 3′SS sequences activated in the absence of Prp18p first focusing on their location relative to the annotated 3′SS and branchpoint sequences (Fig 3a,3b). Plotting the position of alternative 3′SS by distance to the annotated 3′SS revealed that alternative YAG and AAG 3′SS tend to be downstream, while GAG and non-AG sites tend to be upstream of the annotated 3′SS (Fig.3a). The main exceptions to this trend were a group of GAG and BG alternative 3′SS located 2 and 1 nucleotides downstream of the annotated one (Fig.3a). This observation is consistent with the model that spliceosomes lacking Prp18p may exhibit ‘slippage’ when suboptimal 3′SS sequences are located immediately adjacent to the normal 3′SS.

**Figure 3.**
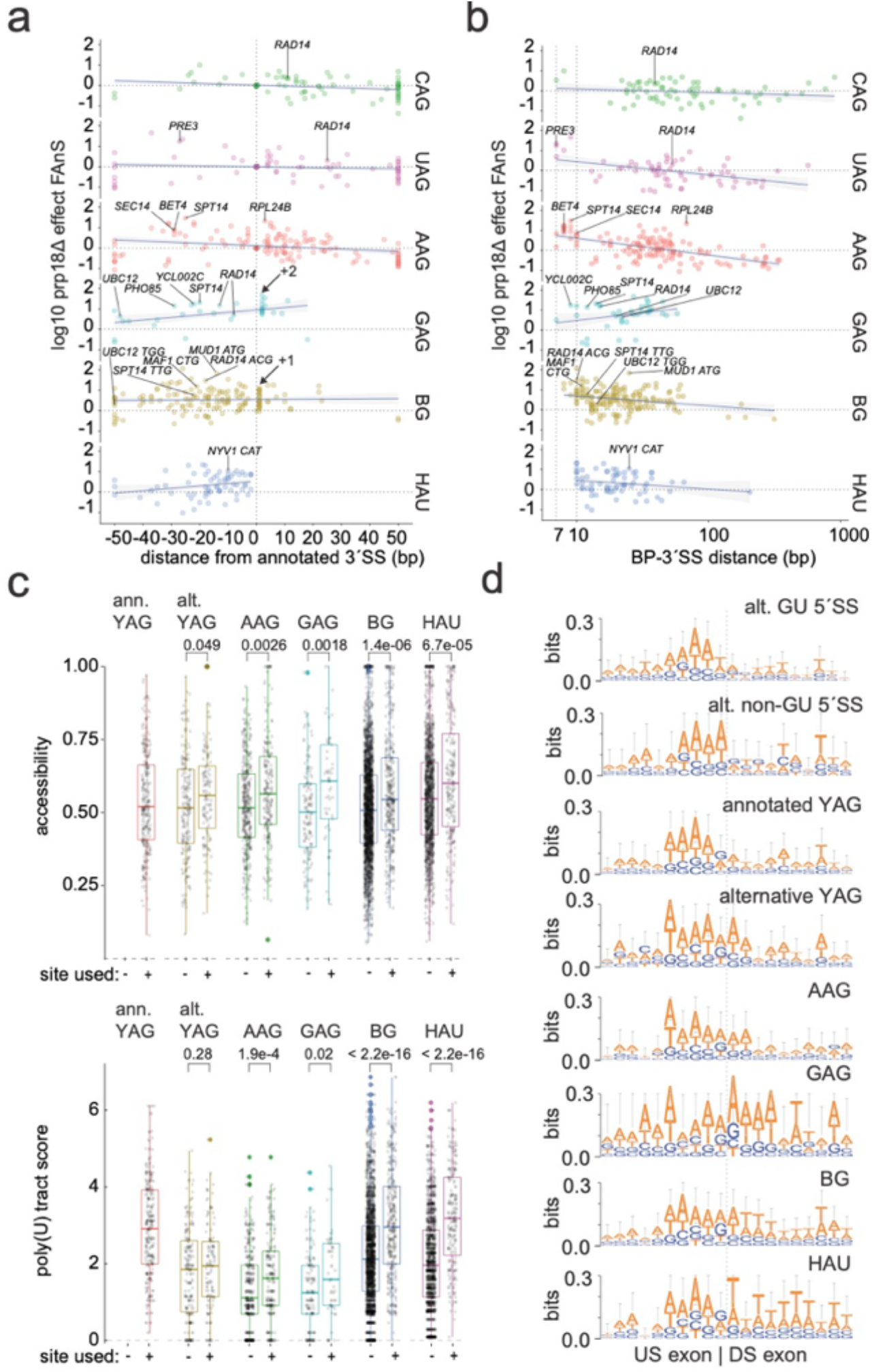
Adjacent sequences and structural contexts modulate alternative 3′SS usage. (a) *prp18*Δ effect (*prp18*Δ*upf1*Δ SE/ *upf1*Δ SE) plotted by the alternative to annotated 3′SS distances for each motif class, where negative numbers represent upstream alt.3′SS. The horizontal line denotes neutral effect of Prp18p. The indicated +1 and +2 positions from the annotated site denote positions enriched for BG and GAG non-canonical splicing motifs, respectively. (b) *prp18*Δ effect as in defined in (a) but plotted by the distance from the branchpoint (BP) to the 3′SS. Note the log scale for the x-axis. (c) Boxplots of 3′SS accessibility and poly(U) tract scores separated by whether the 3′SS gives detectable splicing (+) or not (−) in the *prp18*Δ*upf1*Δ strain. The accessibility of each trinucleotide sequence from 8 nt downstream the branchpoint to 50 nt downstream of the annotated site was calculated as the averaged probability of each base in the 3′SS being unpaired (see Methods). (d) Motif enrichment for the 10 nucleotides upstream and downstream of each junction for each indicated 5′SS and 3′SS class. The vertical dotted line denotes the exon-exon junction.

Many alternative 3′SS used in the absence of Prp18p were located less than 10 nt from the annotated branchpoint (Fig.3b). Whereas the shortest known naturally occurring BP-3′SS distance in budding yeast is 10 nt, we detected 25 cases of alternative 3′SS positioned within 10 nt of the branchpoint, with the shortest distance of 7 nt occurring 7 times (*DYN2*, *PRE3*, *NCE101*, *RUB1*, *RPL40B*, *RPS18B*, *RPS27A*; Fig.3b). These sites were utilized at substantially increased levels in the absence of Prp18p, consistent with spliceosome structural studies that predicted that Prp18p acts a molecular ruler imposing a minimum BP-3′SS distance ^7^.

While we observed a weak relationship between alternative 3′SS efficiency and BP-3′SS distance for pseudo-WT spliceosomes (*upf1*Δ), a stronger negative relationship between alternative 3′SS efficiency and BP-3′SS distance emerged in the absence of Prp18p (Fig S5). This effect was strongest for AG sites, likely owing to the increased numbers of sites utilized downstream of the annotated 3′SS (Fig.3a,3b). While YAG sites downstream of the annotated sites tended to be downregulated in the absence of Prp18p, downstream RAG and BG sites were more often upregulated relative to the annotated sites (Fig 3a). This suggests that the increased utilization of non-canonical 3′SS is not entirely due to Prp18p-deficient spliceosomes favoring shorter BP-3′SS distances, but rather indicative of an independent function for Prp18p in YAG selection. Supporting this model, we noticed an enrichment of upregulated NGG sites at +1 and GAG sites at +2 nt downstream of the annotated 3′SS, where the underlined G represents the G-1 of the annotated YAG 3′SS (Fig 3a; NBG are NGG sites at the +1 position due to the invariant G in annotated sites).

Accessibility of 3′SS within the context of RNA secondary structure can play a major role in determining the major 3′SS used (i.e. the annotated site) ^20,21^. To test whether this impacts alternative 3′SS usage globally, we analyzed accessibility of alternative 3′SS motifs (i.e. the probability of the 3′SS not being included in a secondary structure) in a window from 7 nt downstream the BP to 50 nt downstream the annotated 3′SS. We then separated sites in two groups, according to whether they were detected as alternative 3′SS in either strain. Consistently, alternative 3′SS motifs that are utilized tended to be more accessible than those not utilized, and surprisingly, even more accessible than the annotated 3′SS (Fig.3c). Together these data support a model in which 3’SS accessibility influences its utilization by the spliceosome, consistent with previous studies^20^.

### Poly(U) tract strength is a key determinant of non-canonical (non-YAG) 3′SS efficiency

Next we examined the relationship between poly(U) tracts and 3′SS usage, and calculated a poly(U) score based on position weight matrix scores (PWMS) derived from all annotated 3′SS as defined previously^22^. We found that RAG, BG and HAU alternative 3′SS sites that are utilized exhibited substantially greater poly(U) tract scores than non-utilized sites (Fig.3c). This impact of poly(U) sequence was specific to non-canonical 3′SS sequences, as it was not detected for alternative YAG sites (Fig 3c). This observation suggests that activation of unconventional 3′SS sequences requires the presence of strong poly(U) tract upstream. As alternative 3′SS are in competition with annotated ones, we examined how poly(U) tract scores at the annotated sites impact the splicing efficiency (SE) at alternative 3′SS. Strikingly we observed a significant negative correlation between the poly(U) tract scores of annotated 3′SS and SE of alternative 3′SS (Fig S6a). This was true even when analyzing RPG and non-RPGs separately to account for the observation that RPGs tend to have much stronger poly(U) tracts (Fig S6a). Interestingly, BG and HAU alternative 3′SS exhibited greater poly(U) tract scores relative to the annotated sites in RPG vs non-RPGs (Fig.S6b), indicating greater dependency of non-AG sites on a strong poly(U) tract in the context of RPG splicing (Fig S6b). These differences are significant in light of the fact that GAG sites tend to have weaker poly(U) tracts relative to BG and HAU despite also being predominantly upstream of annotated sites (Fig.3c).

### Exonic adenosines determine the strength of alternative splice sites

Previous work has shown that adenosines in both exons can promote the efficiency of the second step via interactions with the uridine-rich loop 1 of U5 snRNA, and that these interactions are particularly important in the absence of fully functional Prp18p^10,11^. Consistent with these targeted analyses, sequence logo analysis of the alternative 5′SS indicated an enrichment of adenosines in the first 4-5 nt upstream, similar to that observed at annotated 5′SS (Fig.3d). While annotated 3′SS did not show any enrichment for adenosine in the downstream exons, alternative AG 3′SS showed enrichment for A at the +1 and +2 positions, with GAG showing the strongest enrichment from +1 to +4 of the pseudo-exon2 sequences (Fig.3d). The enrichment for U downstream of BG and HAU is likely due in part to the presence of poly(U) tracts upstream of the annotated site, as the majority of BG and HAU sites are found within 20 nt upstream of the annotated site, thus positioned very closely upstream from the poly(U) tract regions preceding the annotated 3′SS. Instead, GAG sites exhibit stronger enrichment of downstream pseudo-exonic adenosines, showing that different non-canonical motifs may rely on different features for efficient usage by the spliceosome (Fig.3d). Overall, these results suggest that the utilization of non-canonical sites requires a combination of intronic and pseudo-exonic features to compete effectively with the annotated sites.

### Mutations in the conserved loop and Helix2 of Prp18p impact splicing fidelity without affecting cellular fitness

*S.cerevisiae* Prp18p contains five α-helices interspersed with short loop regions, including a loop connecting helices 4 and 5 that contains a highly conserved region (CR) of 25 amino acids (Fig.4a; Fig.S7)^23,24^. Cryo-EM structure of the post-catalytic spliceosome revealed that the Prp18p α-helical domain interfaces with the RNase H-like domain of Prp8p at the spliceosome periphery while the conserved loop juts into the active site via a tunnel through Prp8p^7^. This arrangement is proposed to stabilize the core spliceosome, while the Prp18p CR along with the α-finger domain of Prp8p forms a channel within the active site through which the 3′SS can be positioned for exon ligation (Fig 4a). These observations suggest that the CR loop of Prp18p might promote the fidelity of the second step of splicing through its interaction with the 3′SS region. To test this hypothesis, we analyzed the impact of three mutations that were previously reported to disrupt structural domains of Prp18 (Fig S7) (i) a modified deletion of the conserved loop that disrupts growth above 34°C but binds Slu7p efficiently (*prp18-*Δ*CRt*, Δ(S187-I211)); (ii) a double point mutation in helix 2 that inhibits binding to Slu7p (*prp18-h2*, R151E R152E); and (iii) a quadruple point mutation of four conserved residues in helix 5 that severely impacts growth at 30°C but retains the interaction with Slu7p (*prp18-h5*, D223K E224A K234A R235E)^25^. We then analyzed the impact of these mutations on the splicing of two pre-mRNAs that exhibited increased usage of non-canonical 3′SS in the absence of Prp18p: *NYV1* (CAU 3′SS), and *MUD1* (AUG 3′SS).

**Figure 4.**
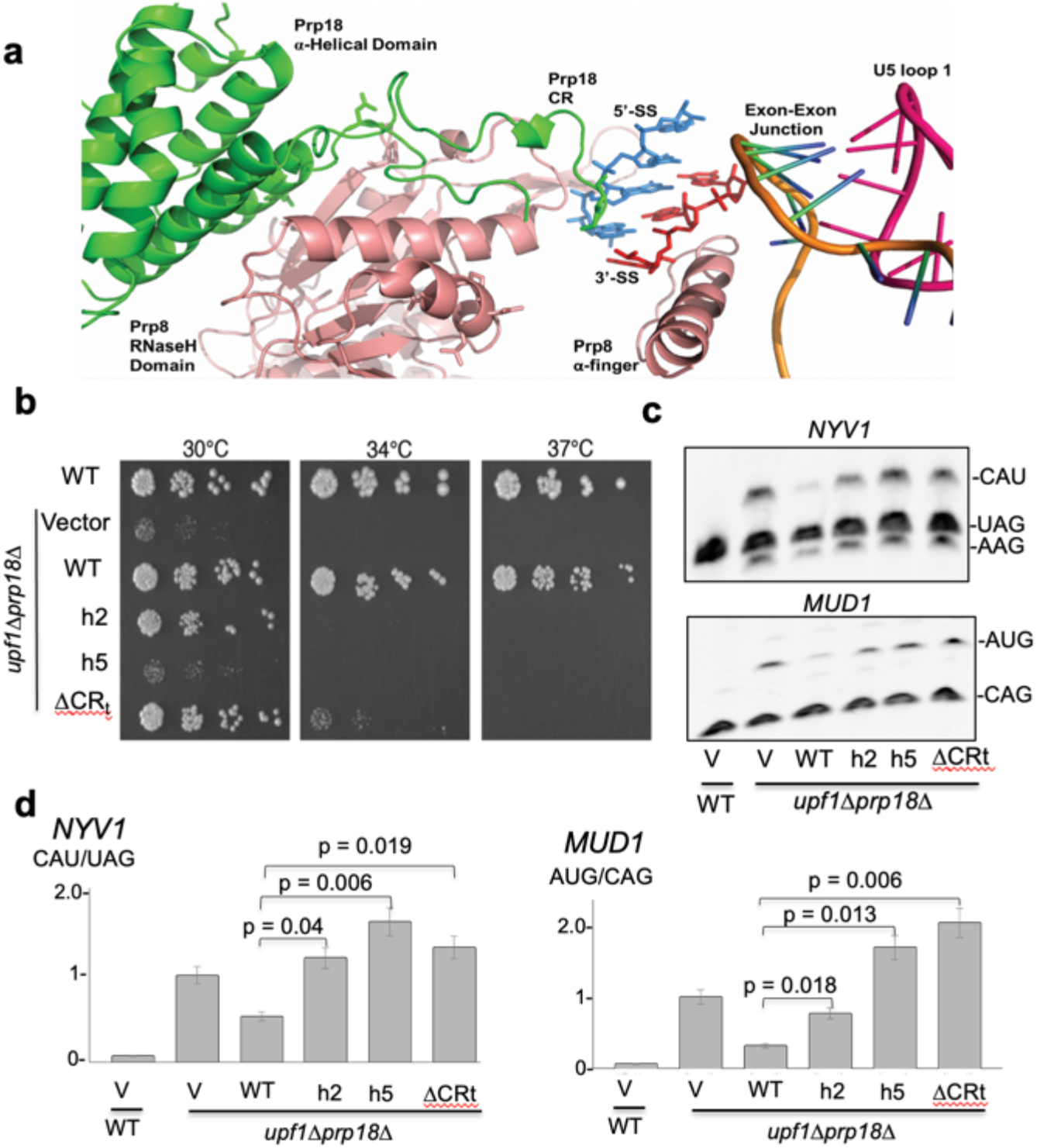
**(a)** Structure of the active site of the post-catalytic spliceosomewith domains of Prp18p (green), Prp8p (salmon), the 5’-SS (blue), the 3’-SS (red), the ligated exons (orange/blue/green), and the loop 1 of the U5 snRNA (pink). **b**. Spot dilution growth assay for wild-type and *prp18*Δ*upf1*Δ strains complemented with various *PRP18* plasmids: empty pUG35 (vector) or derivatives expressing wild-type *PRP18*, a helix 2 mutant (*prp18-h2*), a helix 5 mutant (*prp18-h5*), or a mutant in which the conserved loop is completely deleted (*prp18-*ΔCRt). Strains were grown to exponential phase in CSM-URA media then serially diluted in 3-fold spot dilutions onto CSM-LEU plates and incubated at the indicated temperatures for 3 days. **(c)** RT-PCR analysis of *NYV1* and *MUD1* alternative splicing in wild-type and *prp18*Δ*upf1*Δ strains complemented with an empty vector or plasmids expressing wild-type *PRP18*, a helix 2 mutant (*prp18-h2*), a helix 5 mutant (*prp18-h5*), or a mutant in which the conserved loop is completely deleted (*prp18-*ΔCRt). **(d)** quantification of the usage of the major alternative 3′SS of *NYV1* and *MUD1* relative to the main 3′SS in arbitrary units. The y-axis indicates the ratio of intensities of the bands for the non-canonical site vs. the canonical site, normalized to 1 for the *prp18*Δ*upf1*Δ strain. Error bars represent the standard deviation for three independent biological replicates. P-values are based on T-test analysis (see methods).

Consistent with previous observations, the *prp18-h5* mutant displayed a severe growth defect at 30°C while the growth of the other two mutants was unaffected at this temperature (Fig 4b). This result indicates that the integrity of helix 5 is essential to achieve full functionality of Prp18p, including maintaining normal cell growth. All three mutants showed growth inhibition at 34 and 37°C (Fig.4b), in agreement with previous results^25^. Surprisingly, we found statistically significant increased usage of the non-canonical 3′SS of *NYV1* and *MUD1* in all these mutants at 30°C (Fig.4c, 4d), despite of the fact that only the null and *prp18-h5* mutants display growth defects at this temperature (Fig.4b). Strikingly, usage of the *MUD1* AUG alternative 3′SS is even more pronounced in strains expressing the h5 and the Δ*CRt* mutants than in the *prp18*-null background (Fig.4d), suggesting that these mutants may have dominant phenotypes on the mechanism of 3′SS selection. We conclude that helix mutations of Prp18p can compromise the fidelity of 3′SS selection without necessarily impacting cellular fitness, providing a first line of evidence that the function of Prp18p functions in splicing fidelity and splicing efficiency can be genetically uncoupled.

### Slu7p is necessary to recruit Prp18p but not sufficient to fulfill its 3′SS fidelity functions

Some of the Prp18p mutations that we analyzed above impacted the interaction of Prp18p with Slu7p, so we next asked if Slu7p may influence the ability of Prp18p to promote 3′SS fidelity, as Slu7p has been previously shown to influence 3′SS choice^26^. Prior work has shown that an excess of Slu7p can restore the splicing defect of yeast splicing extracts depleted for Prp18p ^27^. To test whether Slu7p can compensate for the loss of 3′SS fidelity detected in the absence of Prp18p *in vivo*, we overexpressed Slu7p in the *prp18*Δ mutant and assayed growth and splicing phenotypes. Overexpression of Slu7p conferred a partial rescue of the growth defect observed in the *prp18*Δ*upf1*Δ strain (Fig.5a), consistent with previous observations^25^. However, overexpression of Slu7p in the *prp18*Δ*upf1*Δ strain had no effect on the splicing of the *NYV1* mRNA, at the annotated or alternative CAU or AAG sites (Fig.5b). This result suggests that overexpressing Slu7p might rescue growth defects through enhanced splicing of a subset of intron-containing transcripts rather than by improving 3′SS fidelity. This result also reinforces the idea that the function of Prp18p in 3′SS fidelity is separable from its role in promoting splicing efficiency and growth.

We next asked whether the recruitment of Prp18p by Slu7p is required to fulfill Prp18p’s function in the fidelity of 3′SS selection. We analyzed *NYV1* and *MUD1* splicing in the previously described *slu7-11* (H75R, R243G, D267G), *slu7-14* (R173G, W210R, L297Q), and *slu7-EIE* (E215A-I216A-E217A) temperature-sensitive mutant strains ^28^. We found that the *slu7-EIE* allele is the only *slu7* mutant which exhibited significant usage of the non-canonical alternative 3′SS of *NYV1* or *MUD1* at 30°C (Fig.5c, 5d) or 37°C (Fig.5d). This result is consistent with previous data showing that the *slu7-EIE* mutation abolishes the interaction of Slu7p with Prp18p^29^.and therefore prohibits the recruitment of Prp18p to the spliceosome, phenocopying the absence of Prp18p. We conclude that the recruitment of Prp18p by Slu7p is critical to promote the role of Prp18p in the fidelity of 3′SS selection, and that this role cannot be fulfilled by Slu7p alone.

**Figure 5.**
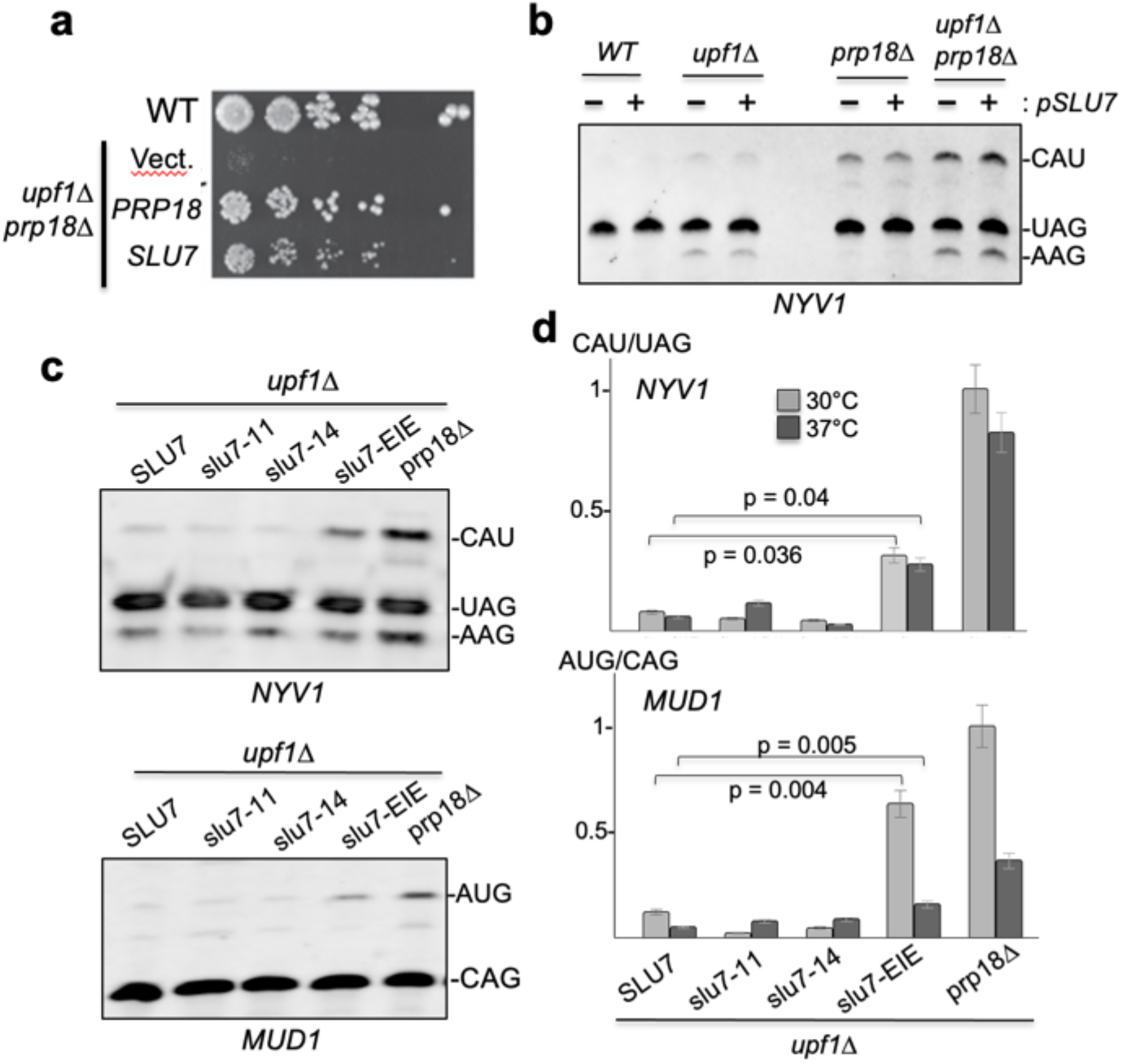
Slu7p is necessary to recruit Prp18p but not sufficient to fulfill its 3′SS fidelity functions. **(a)** Spot dilution growth assay for wild-type and *prp18*Δ*upf1*Δ strain complemented with Yep24 plasmids expressing *PRP18* or overexpression *SLU7* or empty Yep24 (vector) **(b)** RT-PCR analysis of NYV1 alternative 3′SS splicing in WT,*upf1*Δ, *prp18*Δ, and *prp18*Δ*upf1*Δ strains transformed with a vector (−) or a plasmid overexpressing SLU7 (+). **(c)** RT-PCR analysis of *NYV1* and *MUD1* alternative splicing in various slu7 mutants. All strains are in a *upf1*Δ background. **(d)** Quantification of the ratio of usage of the major alternative 3′SS of *NYV1* and *MUD1* relative to the main 3′SS in the various slu7 mutants. Legends as in Fig.4.

### Mutations at the Prp8p • Prp18p interface disable Prp18p fidelity function

Recent structural studies of yeast spliceosomes in various late stages of the spliceosome cycle have shown that Prp18p and Prp8p interact through the RNase H-like domain of Prp8p (Fig. 4a) ^7,30^. Prp8p is the largest spliceosomal protein, a central component of the U5-U4/U6 tri-snRNP and is required for both steps of splicing ^31,32^. Two *PRP8* mutant alleles, *prp8-121* and *prp8-123* were previously shown to confer increased splicing of reporter transcripts carrying point mutations within the 3′SS YAG motif^33^. However, neither of these *prp8* mutants triggered alternative splicing at the non-canonical *NYV1* CAU 3′SS (Fig.S8). While it remains possible that these *prp8* mutations may affect splicing fidelity for other endogenous transcripts, this result demonstrates that the role of Prp18p in promoting 3′SS fidelity is likely to be distinct from the fidelity mechanism that is inhibited in the *prp8-121* and *prp8-12* alleles.

We next tested whether the function of Prp18p in the fidelity of 3′SS selection is promoted by its interactions with Prp8p, and hypothesized that positioning of Prp18p by the Prp8p RNase H domain in the spliceosome active site would be critical to achieve fidelity. We introduced two mutations into the endogenous *PRP8* gene at P1984 and L1988 whose backbone amides are positioned for hydrogen bonding with the side chains of residues N190 and Q181 of Prp18p, respectively, in both the pre-catalytic and post-catalytic spliceosomes (Fig 6a). The *prp8-P1984A-L1988A* (PALA) exhibited significantly increased usage of the alternative 3′-SS of *NYV1* and of *MUD1* (Fig.6b,6c).

**Figure 6.**
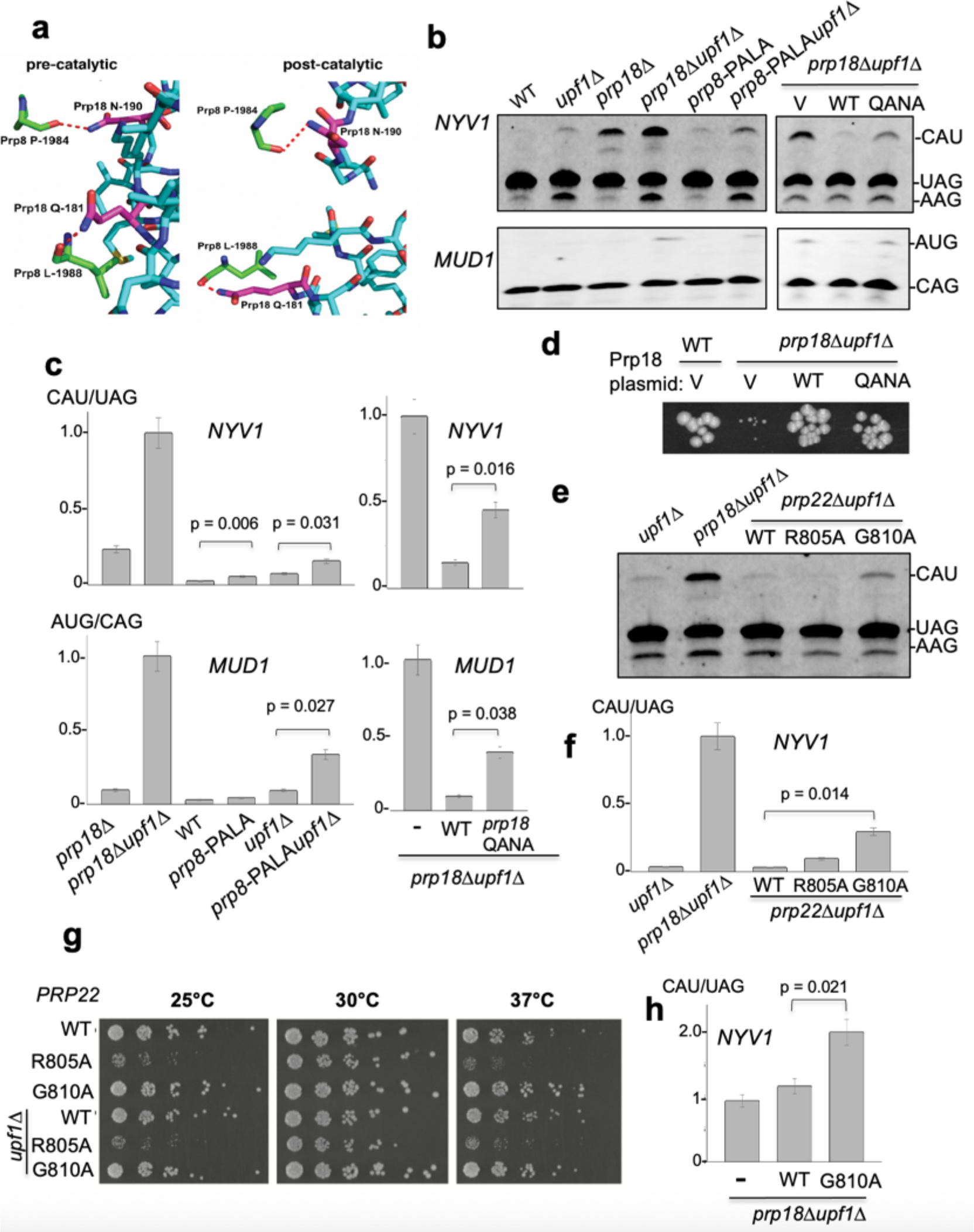
Prp8p and Prp2p22p synergize with Prp18p to promote 3′SS fidelity. **(a)**. Structural details of catalytically activated step II spliceosome ^34^, left, and post-catalytic spliceosome ^7^, right showing interactions between Prp8p P-1984 and Prp18p N-190, and Prp8p L-1988 and Prp18p Q-181. **(b)** RT-PCR analysis of *NYV1* and *MUD1* alternative splicing in a *prp8-PALA* mutant with a double point mutation P1984A and L1988A in WT or *upf1*Δ background. WT, *upf1*Δ, *prp18*Δ and *prp18*Δ *upf1*Δ strains were included for reference. **(c)** Quantification of the ratio of usage of the major alternative 3′SS of *NYV1* and *MUD1* relative to the main 3′SS in the *prp8*-PALA mutant and other mutants analyzed in (b). Legends as in Fig.4. **(d)**. Growth analysis of WT and *upf1*Δ*prp18*Δ transformed with a pUG35 vector (V) or a pUG35 plasmid containing WT *PRP18* (WT) or a double mutant containing a Q181AN191A double mutations (QANA). Strains were grown for 3 days at 30°C. RT-PCR analysis of NYV1 splicing in wild-type, *prp18*Δ, or *prp8* point mutant strains. **(e)** RT-PCR analysis of *NYV1* alternative splicing in wild-type, *prp18*Δ, or *prp18* point mutant strains. All strains were in a *upf1*Δ background. **(f)** Quantification of the ratio of usage of the major alternative 3′SS of *NYV1* for the strains analyzed in (e). Legends as in Fig.4. **(g)** Spot dilution growth assay showing *prp22*Δ null mutant yeast in either a wild-type (top three rows) or *upf1*Δ NMD mutant background (bottom three rows) carrying pRS315 plasmids expressing either wild-type *PRP22*, *prp22-R805A*, or *prp22-G810A* alleles. Strains were grown to exponential phase in CSM-LEU media then serially diluted in 3-fold spot dilutions onto CSM-LEU plates and incubated at the indicated temperatures for 3 days. **(h)** Quantification of the ratio of usage of the major alternative 3′SS of *NYV1* for a *prp18*Δ*upf1*Δ strain transformed with an empty vector (V) or a pRS315 plasmid expressing *PRP22* (WT) or *prp22-G810A*. Legends as in Fig.4.

To further evaluate the importance of the Prp8•Prp18p interface in 3′SS fidelity, we mutated the other side of the interface with two alanine substitutions at residues Q181 and N190 of Prp18p (*prp18*-*QANA* mutant). Strikingly, the *prp18-QANA* allele exhibited a strong activation of the *NYV1* and *MUD1* alternative 3′SS (Fig.6b,6c). However, the QANA allele fully rescued the severe growth defect of the *prp18*Δ*upf1*Δ double mutant (Fig.6d), further demonstrating that the fidelity function of Prp18p can be genetically uncoupled from its role in promoting efficient splicing and general growth. We conclude that the Prp18•Prp8 interface is critical for the fidelity of 3′SS recognition by the spliceosome and to prevent usage of non-canonical 3′splice sites by the spliceosome.

### Prp22p synergizes with Prp18p to prevent usage of non-canonical 3′ splice sites

We next explored whether other splicing factors may assist or synergize Prp18p function in promoting 3′SS fidelity. We began with Prp17p, which stabilizes the C* complex following remodeling by Prp16 but preceding exon ligation^30^, and Cwc21p, which has also been shown to influence 3′SS selection under certain conditions^35^. RT-PCR analysis of *prp17*Δ and *cwc21*Δ null mutants revealed no increased splicing at the non-AG 3′SS of *NYV1* (Fig.S9). We next analyzed mutants of the RNA helicase Prp22p due to its known role in proofreading splicing^9^ and mediating alternative 3′ SS selection^8^. Previous work found increased usage of a non-consensus RAG 3′SS in splicing reporters in two *prp22* mutants harboring mutations within the conserved motif VI, *prp22-R805A* and *prp22-G810A*^9,36^. Strikingly, the *prp22-G810A* allele exhibited a significant accumulation of the *NYV1* CAU alt.3′SS when this mutant was grown at the permissive temperature of 30°C (Fig.6e, 6f). By contrast, the *prp22-R805A* mutant did not appear to impact *NYV1* splicing fidelity (Fig.6e, 6f) despite having much stronger growth defects at reduced and elevated temperatures relative to *prp22-*G810A (Fig.6e, 6f,6g). The growth of these two mutants at different temperatures was consistent with previous reports ^36^(Fig.6g), and inactivation of *UPF1* had not further effect on cellular growth of these mutants (Fig.6g). The observation that the *prp22-G810A* mutant exhibited reduced 3′SS fidelity in conditions where no growth phenotypes are observed (30°C; Fig.6g) strengthens the concept developed from previous experiments with other mutants that splicing fidelity can be uncoupled from optimal cellular growth.

Overall, the finding that alternatively spliced *NYV1* transcripts using the non-canonical 3′SS can be detected in both Prp18p and Prp22p mutants suggests that the two proteins may cooperate in promoting the fidelity of 3′SS selection, with Prp18p playing a primary role in selecting canonical 3′SS sequences and Prp22p playing a downstream role in proofreading as suggested previously ^9^. To test this hypothesis, we expressed the *prp22-G810A* mutant in the *upf1*Δ*prp18*Δ background, with the idea that expressing this mutant should be epistatic to the absence of Prp18p if the two proteins are involved in the same step. Strikingly, expressing the *prp22-G810A* mutant in the *upf1*Δ*prp18*Δ background resulted in an additive effect on the accumulation of the *NYV1* CAU alt.3′SS (Fig 6h) demonstrating that Prp18p and Prp22 function independently to promote the fidelity of 3′SS selection. Our model suggests that Prp18p enhances splicing of canonical 3′SS sequences at a minimum distance of 10 bp from the branchpoint, while Prp22p assumes a downstream proofreading role to reject aberrantly spliced transcripts, possibly because splicing at these non-canonical 3′SS is too slow which would consistent with the classical kinetic proofreading mechanism proposed for splicing helicases^37^.

## Discussion

In this study, we show that Prp18p plays a global role in promoting the selection of canonical 3′SS sequences by the spliceosome. The relaxed fidelity of the spliceosome towards non-canonical 3′ SS sequences is mostly specific to the loss of Prp18p, as it is not detected in many other splicing mutants directly implicated in the 2^nd^ splicing step, including *prp17* and some *prp8* mutants previously described to affect 3′SS fidelity^33^. However, relaxed fidelity is also triggered by mutations that prevent recruitment of Prp18p (such as *slu7-EIE*), or perturb its positioning in the spliceosome active site (such as mutations at the Prp8p-Prp18p interface; Fig.6). Lastly, we demonstrate that the fidelity function of Prp18p is enhanced by the proofreading activity of Prp22p, as shown by the additive effects of inactivating Prp18p and expressing dominant negative versions of Prp22p. Therefore, 3′SS fidelity is a multi-step process in which Prp18p acts to promote fidelity of selection of canonical 3′SS sequences, while Prp22p proofreads splicing variants that use suboptimal 3′SS. The ability of Prp22p to proofread these splicing events might be linked to slower kinetics for the 2^nd^ splicing step of these suboptimal substrates consistent with the kinetic proofreading model proposed for splicing helicases^37^.

Alternative, non-consensus 3′SS used in the absence of Prp18p are highly diverse in their sequence, which raises the question why specific sites are being used as opposed to others which are very similar and located nearby. We show here that several intronic and pseudo-exonic features such as the distance from the branchpoint to the 3′SS, the presence of poly(U) tracts upstream the alternative and annotated sites, adenosine content in the exons or pseudo-exons, and accessibility of the site within RNA secondary structure likely combine to dictate the efficiency of alternative 3′SS usage. Interestingly, different budding yeast species have been shown to exhibit different ranges of BP-3′SS distances, with the salt-tolerant species *Debaryomyces hansenii* exhibiting the shortest predicted BP-3′SS distances of predominantly 7-8 nt^38^. The observation that Prp18p appears to impose a minimum BP-3′ SS distance suggests that inter-species differences in the Prp18p protein sequence might set a lower limit on the allowable BP-3′ SS distance.

After the first chemical step of splicing, the spliceosome needs to thread the sequence between the branchpoint and the 3′SS to insert the 3′SS in the active site. This threading process might be slower in the absence of Prp18p, which would explain why non-canonical 3′SS upstream from the normal 3′SS are used more efficiently (Fig.3a,b). We propose a model wherein in the absence of Prp18p, non-canonical 3′SS sequences are more likely to engage in promiscuous interactions with the first intronic nucleotide à la Parker and Siliciano ^39^ and position the 3′SS for ligation as shown in spliceosome structures ^7^ (Fig S10a). In light of the global activation of HAU alternative 3′SS sequences, we propose that in the absence of Prp18p, there may be increased flexibility in the active site which allows for an alternative base-pairing with the 3′SS U**_-1_** and the 5′SS G**_+1_**(Fig.S10b). Substituting a cytosine or adenosine at the last position of the 3′SS would introduce a steric clash or incompatible hydrogen bond donor/acceptor arrangements that would disrupt the non-Watson Crick interaction with G1 (Fig S10c,d), providing a rationale for the preference for Us we observe at the last position of non-G alternative 3′ SS. While the necessity to have a sequence devoid of secondary structure for usage of a particular 3′SS is obvious, the mechanistic basis for the necessity of the poly(U) tract upstream is poorly understood. While polypyrimidine tracts can be recognized by specific splicing factors (eg U2AF65 in higher eukaryotes), it does not explain the specific importance of U over C in the selection process. If the threading model for insertion of the 3′SS in the active site of the spliceosome is correct, the presence of a poly(U) stretch might facilitate threading because uracil is the smallest base, and the absence of an amino group in the base might prevent promiscuous hydrogen bonding with spliceosome components that might potentially slow down the threading process.

This function in the fidelity of 3′SS selection can be genetically uncoupled from its functions in splicing efficiency and general growth, as shown by the distinct effects of *cis*-acting mutants of Prp18p and overexpression of Slu7p. Interestingly, we also observed this uncoupling in the *prp22-G810A* mutant (Fig 6), which exhibits reduced splicing fidelity at a temperature at which splicing efficiency and growth are unaffected. These results are particularly important as they rule out that the loss of fidelity of the spliceosome for consensus sequences is due to a global decrease in splicing efficiency or to an indirect effect of slow growth. The overall observation that point mutations in Prp18p, Slu7p and Prp22p can activate new splice sites without grossly impacting cellular growth (at least in normal growth conditions) suggests that a minor accumulation of aberrantly spliced products is not a major deterrent to cellular fitness. This suggests a possible mechanism whereby new splice isoforms could be explored during evolution.

## Materials and Methods

### Yeast strain and plasmid construction

Yeast strain construction, Northern blot analysis, RT-PCR analysis, molecular cloning, restriction enzyme-based plasmid construction, site-directed mutagenesis, and spot dilution assays were performed as previously described ^40,41,42^. Construction of the PRP18/pUG35 and PRP18/YEp24 base plasmids was accomplished using Gibson assembly^43^ (New England Biolabs #E2611). Unless otherwise indicated in the figure legends, all strains were derived from the BY4741 background. The *PRP8* genomic mutations were introduced using the CRISPR-Cas system as previously described^44^. The *SLU7* and *PRP22*/*PRP43* mutant strains and plasmids were generously provided by Beate Schwer (Cornell U.) and Jonathan Staley (U. Chicago), respectively.

### Yeast growth, RNA preparation and sequencing

Yeast strains were grown and harvested as previously described ^40,41^. For heat-sensitive mutant strains, cell cultures were first grown to exponential phase at 25°C, a portion harvested, and the remaining culture shifted to 37° and incubated at that temperature for 1 hour. For cold-sensitive mutant strains, cell cultures were first grown to exponential phase at 30°C, a portion harvested, and the remaining culture split and shifted to 16°C or 25°C and incubated at those temperatures for 2 hours. For the RNA-Seq analysis, *upf1Δ∷HIS3MX6* and *upf1Δ∷HIS3MX prp18Δ∷kanMX6* colonies were inoculated for overnight growth in YPD, seeded in 50 mL fresh YPD the following day at OD_600 nm_ 0.05 and grown to mid-log phase OD_600 nm_ of 0.6. Samples were harvested by spinning down at 3k rpm for 5 minutes, flash-frozen in liquid nitrogen, and total RNA was purified by standard phenol-chloroform extraction as described previously^41^. Fragmented RNA seq libraries were prepared using the Illumina TruSeq stranded mRNA library prep kit. 2 × 100bp sequencing on the HiSeq2000 and subsequent demultiplexing were performed by Macrogen/Axeq (South Korea).

### Computational methods

A detailed description of all computational steps is outlined in the Supplemental Methods. All scripts for the Comparison of Multiple alignment Programs for Alternative Splice Site discovery (COMPASS) read processing pipeline are available on GitHub (https://github.com/k-roy/COMPASS). COMPASS includes the following steps: pre-processing of raw reads, alignment with multiple aligners, post-alignment processing, selection of the best scoring alignment for each read, and quality filtering of junctions. The aligners STAR^16^ and BBMap [https://sourceforge.net/projects/bbmap, https://www.osti.gov/biblio/1241166] were used in this study. Scripts for downstream analyses are available on GitHub (https://github.com/k-roy/COMPASS/tree/master/yeast_Prp18).

### Splicing efficiency (SE) and fraction of annotated splicing (FAnS) calculation

We calculated splicing efficiency (SE) as the ratio of the reads mapping to each splice junction over the sum of all reads mapping to both spliced and unspliced junctions for each intron. Unspliced mRNA harbors both 5′SS and 3′SS junctions so the combined read counts for these junctions are divided by 2. For each alternative splice junction, we calculated a fraction of annotated splicing (FAnS) as the ratio of the reads mapping to the junction relative to the reads mapping to the annotated junction. For both SE and FAnS, we also calculated a *prp18*Δ effect by taking the ratio of the *prp18*Δ *upf1*Δ strain relative to the *upf1*Δ strain.

### Intron sequence and structure prediction analyses

For each 3′ SS, the intronic sequence was examined for the sequence most closely matching the branchpoint consensus TACTA<u>A</u>C, with each mismatch penalized by a score of 1, and an additional penalty of 2 was given if the conserved branched adenosine (underlined) was not present. The sequence with the lowest mismatch score was assigned as the most likely branchpoint, and ties were broken by taking the branchpoint with the shortest distance to the 3′SS. The linear branchpoint (BP)–3′SS distance was defined as the number of nucleotides between the branching A of the BP and the 3′SS, such that TACTA**<u>A</u>** CACNNNN|TAG would be a distance of 10 nt, in line with previous conventions ^20,38^. See COMPASS_analyze_splice_junction_profiles_for_individual_samples.py.

Effective branchpoint (BP)–3′SS distances and RNA tri-nucleotide accessibilities in the BP–3′SS region were calculated similarly to a previous report ^20^. For each annotated and alternative 3′SS, four windows of increasing length size with lengths n+5, n+10, n+15, and n+20 respectively, where n is the distance between the BP and 3′SS, were folded with rnafold version 2.4.6 with options **--partfunc --constraint**^45^. The **--constraint** option enables keeping the branch sequence and 8 nt downstream unpaired as this sequence has been proposed to be kept in an unpaired configuration by the spliceosome^20^. The effective distance from the branchpoint is determined from the lowest energy structure from the parens/bracket notation from rnafold. The effective distance of any base X is calculated by examining the fold in between base X and the branchpoint. The number of bases between base X and the branchpoint which are paired with other bases between base X are subtracted from the linear distance, with bases involved in loops treated as paired. The **--partfunc** option of rnafold calculates the partition function and base pairing probability matrix which are written to dp.ps files. For each base, the probability of being paired with each other base is summed to give a probability of each base being in a base pair. The accessibility of each base was then calculated as 1 minus this probability. The accessibility value for each trinucleotide in the region from 7 nt downstream the branchpoint to 5 nt downstream the splice site was calculated as the average of the accessibility values of the three bases averaged over the four windows. For the comprehensive analysis of each annotated intron where sites are analyzed as used versus unused as shown in Fig 3C, 50 nt were added to the end of the annotated 3′SS. See COMPASS_write_BP_3SS_to_fasta.py and COMPASS_analyze_BP_3SS_RNA_folds_for_annotated_introns.py. The polyU score of each trinucleotide in the region from 7 nt downstream the branchpoint to 5 nt downstream the annotated splice site was calculated using the position weight matrix scores (PWMS) of the yeast polyU tract from a previous study ^22^.

### RT-PCR Analysis and quantification of alternative splice site usage

RT-PCR analysis of alternative splice site usage using fluorescently labeled primers was performed as described^46^. Quantification of the bands corresponding to each isoform was measured using ImageJ 1.53K. To determine the intensity of each band, three separate boxes were created and the average intensity of the three was used as the band intensity. The background intensity for each lane was subtracted from the average band intensity in order to remove background bias. Samples of interest were compared in triplicate using two-tailed t-tests.

## Data availability

All raw and processed data from this study have been submitted to the NCBI Gene Expression Omnibus (GEO; http://www.ncbi.nlm.nih.gov/geo/) under accession number GSE131797.

## Acknowledgements

This work is dedicated to the memory of Christine Guthrie. We thank Sebastian Fica, Jonathan Staley, Beate Schwer and Raymond O’Keefe for insights and/or sharing of materials. This work was supported by grant GM 130370 to GFC. KRR and JG were supported by National Institutes of Health training program T32 GM007185.

## Author Contributions

Conceptualization, JG, KRR, GFC; Methodology, JG, KRR, GFC; Software: KRR; Investigation, JG, KRR, DN, MA, JS, GL; Data Curation, KRR.; Statistical Analysis: KRR, DN; Writing: KRR, JG, GFC; Visualization, JG, KRR, DN; Supervision, JG, GFC; Project Administration: GFC. Funding Acquisition, GFC.

## References

1. Stanley, R. F. & Abdel-Wahab, O. Dysregulation and therapeutic targeting of RNA splicing in cancer. Nat Cancer 3, 536–546 (2022).

2. Wu, S., Romfo, C. M., Nilsen, T. W. & Green, M. R. Functional recognition of the 3’ splice site AG by the splicing factor U2AF35. Nature 402, 832–835 (1999).

3. Schwer, B. & Guthrie, C. A conformational rearrangement in the spliceosome is dependent on PRP16 and ATP hydrolysis. EMBO J 11, 5033–5039 (1992).

4. Wilkinson, M. E., Fica, S. M., Galej, W. P. & Nagai, K. Structural basis for conformational equilibrium of the catalytic spliceosome. Mol Cell 81, 1439–1452.e9 (2021).

5. Ohrt, T. et al. Molecular dissection of step 2 catalysis of yeast pre-mRNA splicing investigated in a purified system. RNA 19, 902–915 (2013).

6. Umen, J. G. & Guthrie, C. Prp16p, Slu7p, and Prp8p interact with the 3’ splice site in two distinct stages during the second catalytic step of pre-mRNA splicing. RNA 1, 584–597 (1995).

7. Wilkinson, M. E. et al. Postcatalytic spliceosome structure reveals mechanism of 3ʹ–splice site selection. Science (1979) 358, 1283–1288 (2017).

8. Semlow, D. R., Blanco, M. R., Walter, N. G. & Staley, J. P. Spliceosomal DEAH-Box ATPases Remodel Pre-mRNA to Activate Alternative Splice Sites. Cell 164, (2016).

9. Mayas, R. M., Maita, H. & Staley, J. P. Exon ligation is proofread by the DExD/H-box ATPase Prp22p. Nat Struct Mol Biol 13, 482–490 (2006).

10. Crotti, L. B., Bacikova, D. & Horowitz, D. S. The Prp18 protein stabilizes the interaction of both exons with the U5 snRNA during the second step of pre-mRNA splicing. Genes Dev 21, 1204–1216 (2007).

11. Crotti, L. B. & Horowitz, D. S. Exon sequences at the splice junctions affect splicing fidelity and alternative splicing. Proc Natl Acad Sci U S A 106, 18954–18959 (2009).

12. Kawashima, T., Douglass, S., Gabunilas, J., Pellegrini, M. & Chanfreau, G. F. Widespread Use of Non-productive Alternative Splice Sites in Saccharomyces cerevisiae. PLoS Genet 10, e1004249 (2014).

13. Kawashima, T., Pellegrini, M. & Chanfreau, G. F. Nonsense-mediated mRNA decay mutes the splicing defects of spliceosome component mutations. RNA 15, 2236–47 (2009).

14. Ares Jr., M., Grate, L. & Pauling, M. H. A handful of intron-containing genes produces the lion’s share of yeast mRNA. RNA 5, 1138–1139 (1999).

15. Pleiss, J. A., Whitworth, G. B., Bergkessel, M. & Guthrie, C. Rapid, transcript-specific changes in splicing in response to environmental stress. Mol Cell 27, 928–937 (2007).

16. Dobin, A. et al. STAR: ultrafast universal RNA-seq aligner. Bioinformatics 29, 15–21 (2013).

17. Aslanzadeh, V., Huang, Y., Sanguinetti, G. & Beggs, J. D. Transcription rate strongly affects splicing fidelity and cotranscriptionality in budding yeast. Genome Res 28, 203–213 (2018).

18. Gould, G. M. et al. Identification of new branch points and unconventional introns in Saccharomyces cerevisiae. RNA 22, 1522–1534 (2016).

19. Schreiber, K., Csaba, G., Haslbeck, M. & Zimmer, R. Alternative Splicing in Next Generation Sequencing Data of Saccharomyces cerevisiae. PLoS One 10, (2015).

20. Meyer, M., Plass, M., Perez-Valle, J., Eyras, E. & Vilardell, J. Deciphering 3’ss selection in the yeast genome reveals an RNA thermosensor that mediates alternative splicing. Mol Cell 43, 1033–1039 (2011).

21. Plass, M., Codony-Servat, C., Ferreira, P. G., Vilardell, J. & Eyras, E. RNA secondary structure mediates alternative 3’ss selection in Saccharomyces cerevisiae. RNA 18, 1103–1115 (2012).

22. Ma, P. & Xia, X. Factors affecting splicing strength of yeast genes. Comp Funct Genomics 2011, (2011).

23. Bacikova, D. & Horowitz, D. S. Genetic and functional interaction of evolutionarily conserved regions of the Prp18 protein and the U5 snRNA. Mol Cell Biol 25, 2107–2116 (2005).

24. Jiang, J., Horowitz, D. S. & Xu, R.-M. Crystal structure of the functional domain of the splicing factor Prp18. Proceedings of the National Academy of Sciences 97, 3022–3027 (2000).

25. Bacikova, D. & Horowitz, D. S. Mutational analysis identifies two separable roles of the Saccharomyces cerevisiae splicing factor Prp18. RNA 8, 1280–1293 (2002).

26. Frank, D. & Guthrie, C. An essential splicing factor, SLU7, mediates 3’ splice site choice in yeast. Genes Dev 6, 2112–2124 (1992).

27. Zhang, X. & Schwer, B. Functional and physical interaction between the yeast splicing factors Slu7 and Prp18. Nucleic Acids Res 25, 2146–2152 (1997).

28. Aronova, A., Bacíková, D., Crotti, L. B., Horowitz, D. S. & Schwer, B. Functional interactions between Prp8, Prp18, Slu7, and U5 snRNA during the second step of pre-mRNA splicing. RNA 13, 1437–1444 (2007).

29. James, S. A., Turner, W. & Schwer, B. How Slu7 and Prp18 cooperate in the second step of yeast pre-mRNA splicing. RNA 8, 1068–1077 (2002).

30. Fica, S. M. et al. Structure of a spliceosome remodelled for exon ligation. Nature 542, 377–380 (2017).

31. Grainger, R. J. & Beggs, J. D. Prp8 protein: At the heart of the spliceosome. RNA 11, 533–557 (2005).

32. Horowitz, D. S. The mechanism of the second step of pre-mRNA splicing. Wiley Interdiscip Rev RNA 3, 331–350 (2012).

33. Umen, J. G. & Guthrie, C. Mutagenesis of the yeast gene PRP8 reveals domains governing the specificity and fidelity of 3’ splice site selection. Genetics 143, 723–739 (1996).

34. Yan, C., Wan, R., Bai, R., Huang, G. & Shi, Y. Structure of a yeast step II catalytically activated spliceosome. Science (1979) 355, (2017).

35. Gautam, A., Grainger, R. J., Vilardell, J., Barrass, J. D. & Beggs, J. D. Cwc21p promotes the second step conformation of the spliceosome and modulates 3ʹ splice site selection. Nucleic Acids Res 43, 3309 (2015).

36. Schwer, B. & Meszaros, T. RNA helicase dynamics in pre-mRNA splicing. EMBO J 19, 6582–6591 (2000).

37. Staley, J. P. & Guthrie, C. Mechanical devices of the spliceosome: motors, clocks, springs, and things. Cell 92, 315–326 (1998).

38. Gahura, O., Hammann, C., Valentová, A., Půta, F. & Folk, P. Secondary structure is required for 3ʹ splice site recognition in yeast. Nucleic Acids Res 39, 9759–9767 (2011).

39. Parker, R. & Siliciano, P. G. Evidence for an essential non-Watson–Crick interaction between the first and last nucleotides of a nuclear pre-mRNA intron. Nature 1993 361:6413 361, 660–662 (1993).

40. Gabunilas, J. & Chanfreau, G. Splicing-Mediated Autoregulation Modulates Rpl22p Expression in Saccharomyces cerevisiae. PLoS Genet 12, e1005999 (2016).

41. Roy, K. & Chanfreau, G. Stress-Induced Nuclear RNA Degradation Pathways Regulate Yeast Bromodomain Factor 2 to Promote Cell Survival. PLoS Genet 10, (2014).

42. Roy, K., Gabunilas, J., Gillespie, A., Ngo, D. & Chanfreau, G. F. Common genomic elements promote transcriptional and DNA replication roadblocks. Genome Res 26, 1363–1375 (2016).

43. Gibson, D. G. et al. Enzymatic assembly of DNA molecules up to several hundred kilobases. Nature Methods 2009 6:5 6, 343–345 (2009).

44. Gillespie, A., Gabunilas, J., Jen, J. C. & Chanfreau, G. F. Mutations of EXOSC3/Rrp40p associated with neurological diseases impact ribosomal RNA processing functions of the exosome in S. cerevisiae. RNA 23, 466–472 (2017).

45. Lorenz, R. et al. ViennaRNA Package 2.0. Algorithms for Molecular Biology 6, (2011).

46. Weathers, I., Gabunilas, J., Samson, J., Roy, K. & Chanfreau, G. F. Protocol for High-Resolution Mapping of Splicing Products and Isoforms by RT-PCR Using Fluorescently Labeled Primers. STAR Protoc 1, (2020).

