## Supplemental Figures and Methods for "Spliceosomal mutations decouple 3′ splice site fidelity from cellular fitness"

**Figure S1. Normalized read counts and splicing efficiency for RPGs versus non-RPGs.** Shown are boxplots depicting normalized mean reads for *upf1*Δ (top) and *prp18*Δ*upf1*Δ (bottom) as a function of annotated and alternative 3´SS and 5´SS.

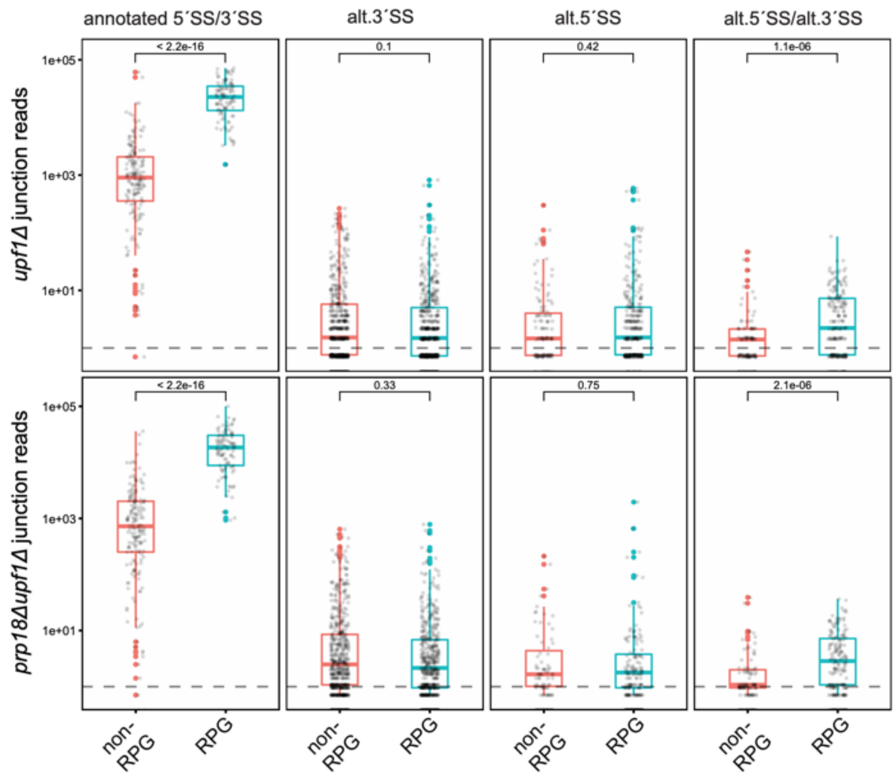


**Figure S2.** The fraction of annotated splicing (FAnS) is shown for the subset of alternative AG 3´SS in the current dataset (x-axis) which were also identified in a recent splicing dataset from budding yeast (y-axis) (Aslanzadeh et al., 2018).

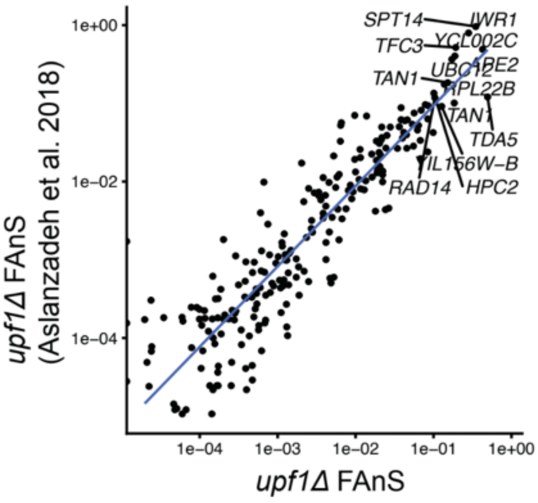

**Figure S3. Impact of Prp18 inactivation on splicing efficiency (SE) and Fraction of Annotated Splicing (FAnS) for alternative junctions.**

**(a)** Boxplot of the *prp18*Δ effect of splicing efficiency (*prp18*Δ*upf1*Δ SE/ *upf1*Δ SE) for alternative junctions**.** Log10 less than zero represents decreased splicing in the absence of Prp18p. Note that while most alternative 5´SS exhibit decreased SE, and on the whole alternative 3´SS have decreased SE, a substantial fraction of alternative 3´SS show increased SE despite the overall negative impact of *prp18*Δ on splicing. **(d)** Boxplot of the *prp18*Δ effect for fraction of annotated splicing (FAnS)

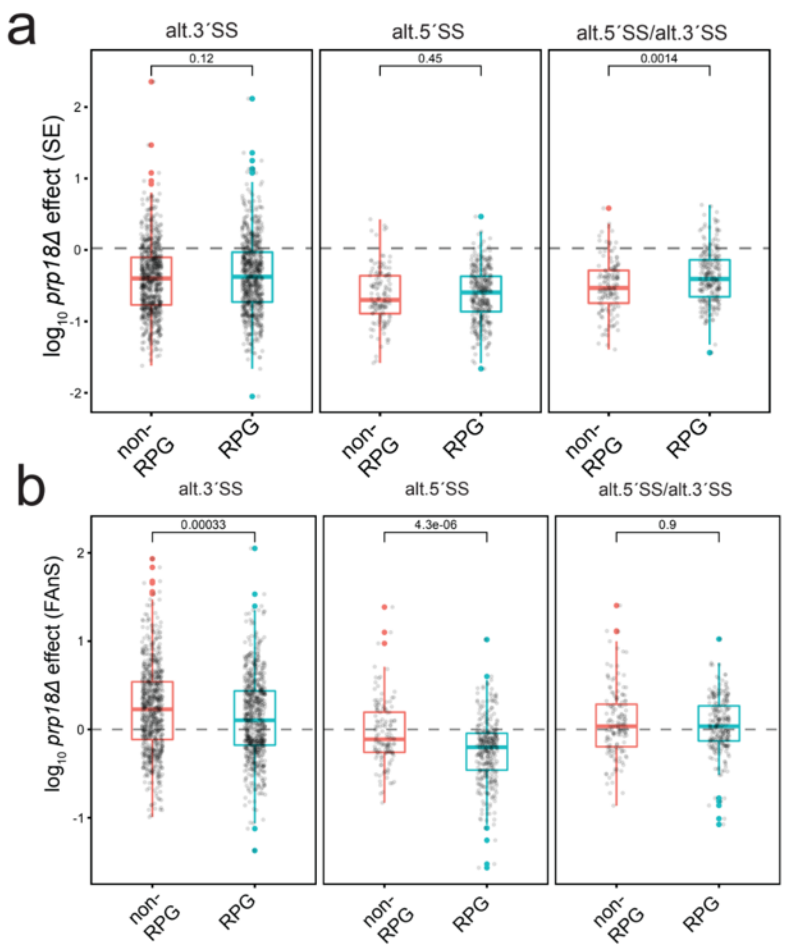

**Figure S4. Splicing efficiency (SE) by motif class in *upf1Δ* vs *prp18Δupf1Δ*. (a)** Log-scale scatter plots of the SE for each splice junction faceted by trinucleotide sequence. Each point corresponds to a distinct splice junction passing the junction quality filters. **(b)** Same as in (a) but grouped by motif class, with each point colored according to whether it harbors an annotated or alternative 5´SS and 3´SS.

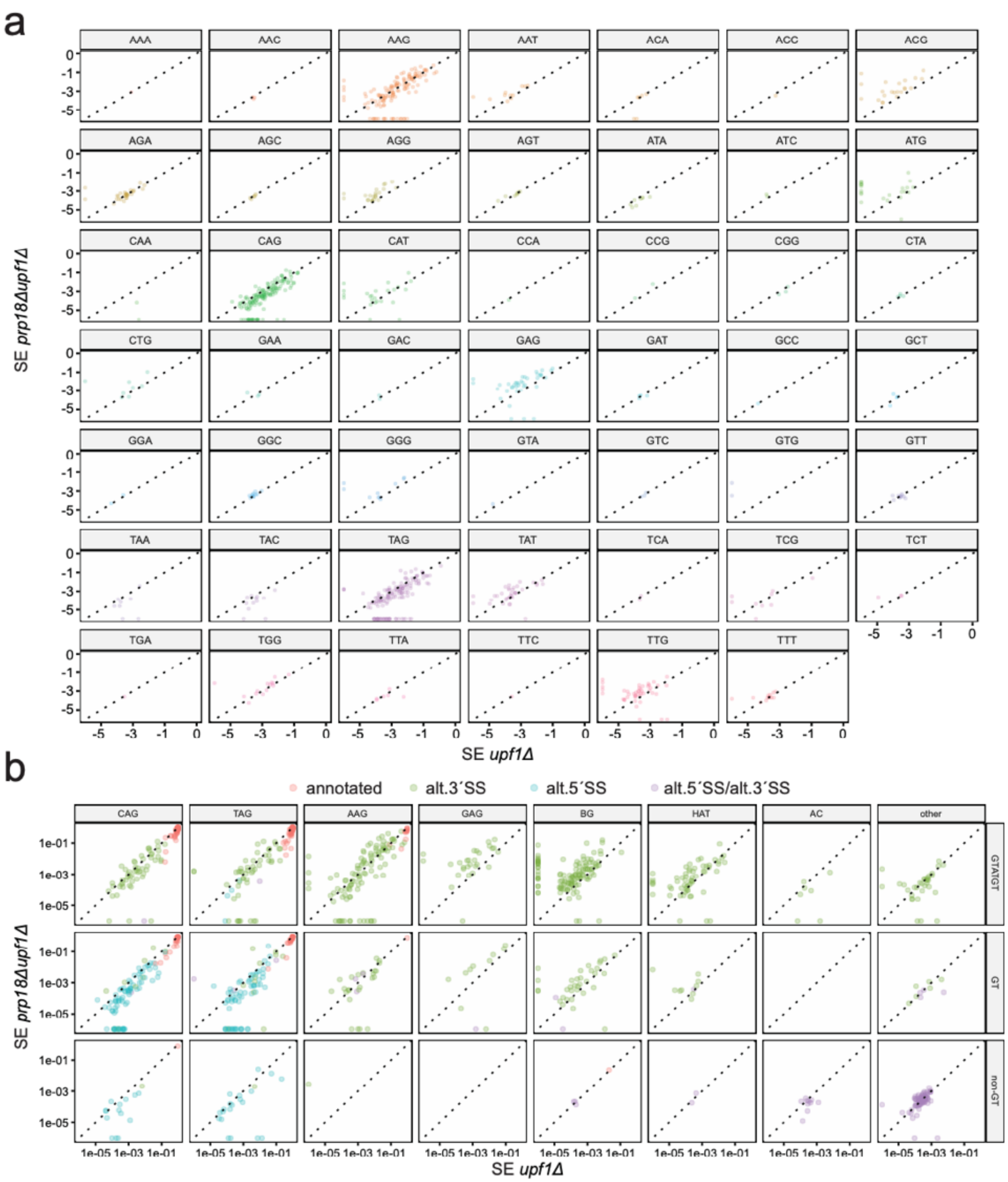

**Figure S5.** Branchpoint (BP) to 3´SS distance distribution and relationship with splicing efficiency. (a) RNA structure between the BP and the 3´SS effectively reduces the physical distance between these two sites. The density distribution for each 3´SS motif is plotted by distance from the branchpoint. BP-3´SS regions for all alternative 3´SS were separated according to whether there was no predicted structure (purple dashed-dotted line), or predicted structure, where the linear distance is shown as a solid turquoise line, and the effective distance shown as a red dotted line. (see Methods for calculation details). (b) Regression analysis on the SE for alt. 3´SS observed in both upf1Δ and prp18Δupf1Δ. The trendline shows a more significant dependence on BP-3´SS distance for spliceosomes lacking Prp18 than for WT spliceosomes.

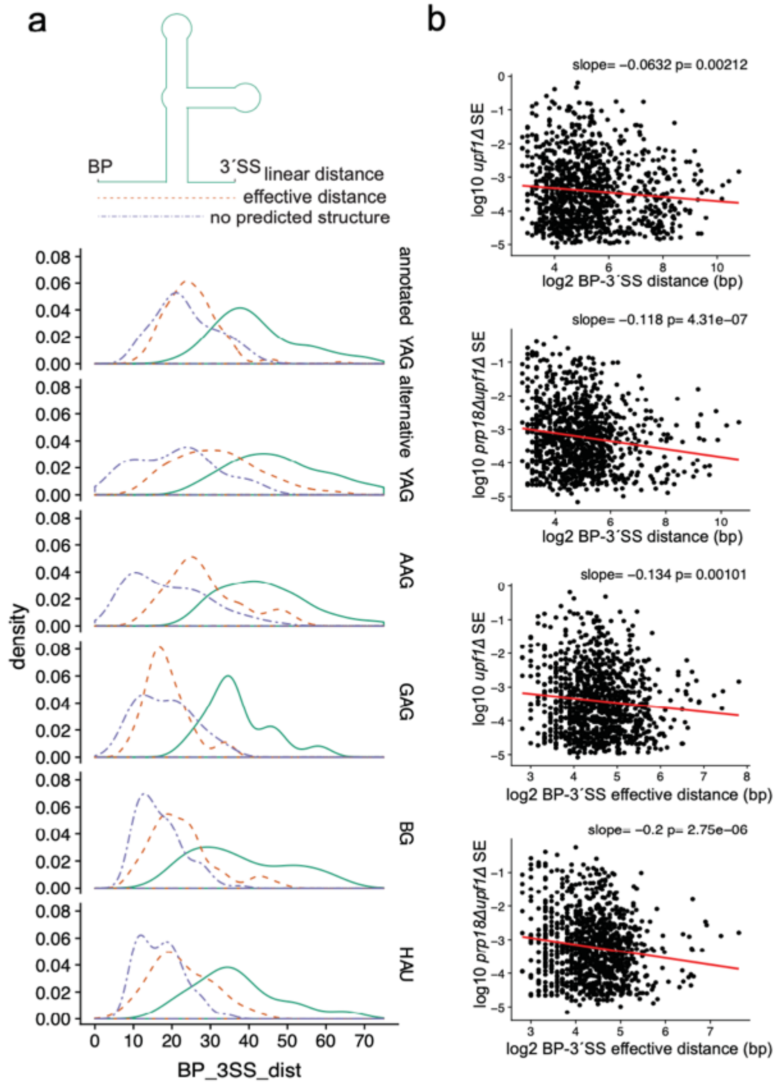

**Figure S6. Impact of Poly(U) content on alternative 3′SS usage.**
(a) Regression analysis of the splicing efficiency for alternative 3′SS in upf1Δ as a function of the poly(U) tract strength of the annotated 3′SS, which is in competition with the alternative sites. The trend lines are separated according to whether the intron belongs to an RPG (red) or non-RPG (blue). (b) Distribution of the difference in U-score between the annotated and alternative sites, according to whether the intron resides in non-RPGs (left) which tend to have lower U-scores, or in RPGs (right), which exhibit higher U-scores.

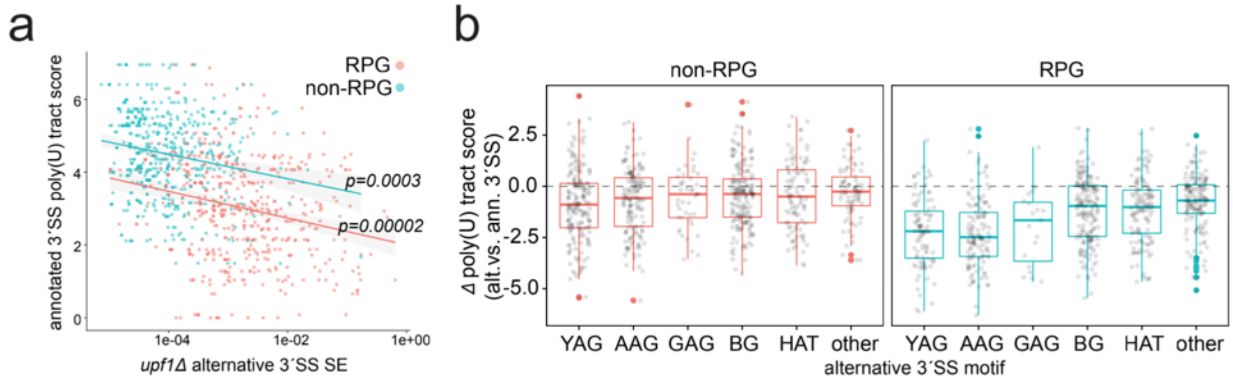


**Figure S7. Summary of Prp18 Conserved region (CR) and Helix 2/5 mutations (a)** Protein sequence of Prp18p with the introduced mutations in red (top). The mutations for each *prp18* allele are summarized in the table at the bottom.

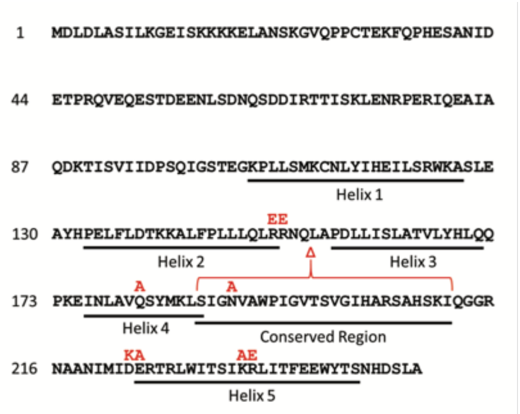


| Name | Mutation(s) |
|---|---|
| ***prp18-h2*** | **R151E R152E** |
| ***prp18-h5*** | **D223K E224A K234A R235E** |
| ***prp18-ΔCRt*** | **Δ(S187-I211)** |
| ***prp18(Q181A/N190A)*** | **Q181A N190A** |

**Figure S8. 3′-SS fidelity mutants of *PRP8* do not activate *NYV1* alternative 3´-SS.**

RT-PCR analysis of of *NYV1* splicing in WT, *prp18*Δ, or *prp8* point mutant strains in either NMD-positive or mutant backgrounds (UPF1 WT / Δ).

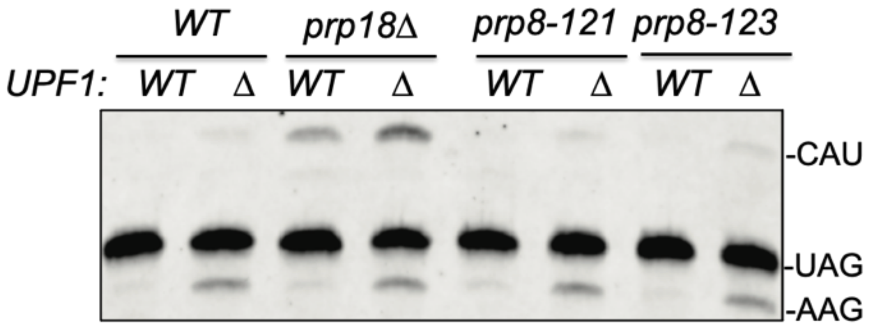


**Figure S9. The *NYV1* alternative 3´-SS is not activated in yeast knockouts of *prp17* or *cwc21*.** RT-PCR analysis of of *NYV1* splicing in WT, *prp18*Δ, *prp17*Δ, *cwc21*Δ and double mutants combined with the *upf1*Δ.

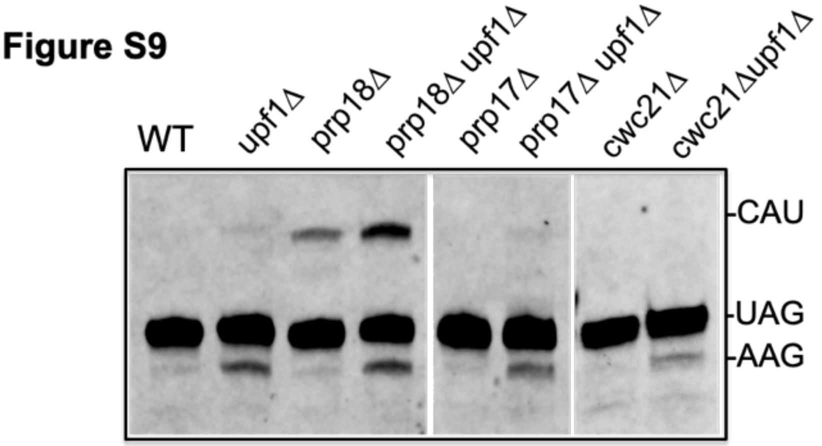

**Figure S10. Non-Watson-Crick base pairs between the first and last intronic nucleotides. (a).** Hydrogen bonding between the intronic G(+1) and G(-1) nucleotides in a non-Watson-Crick base pair. Image created in PyMOL from a structure of the post-catalytic spliceosome published by (Wilkinson et al., 2017). Hypothetical bonding or steric clashes between G(+1) and the following nucleotides in the -1 position: U **(b)**, A **(c)**, or C **(d)**. Yellow curves represent possible steric clashes between molecules.

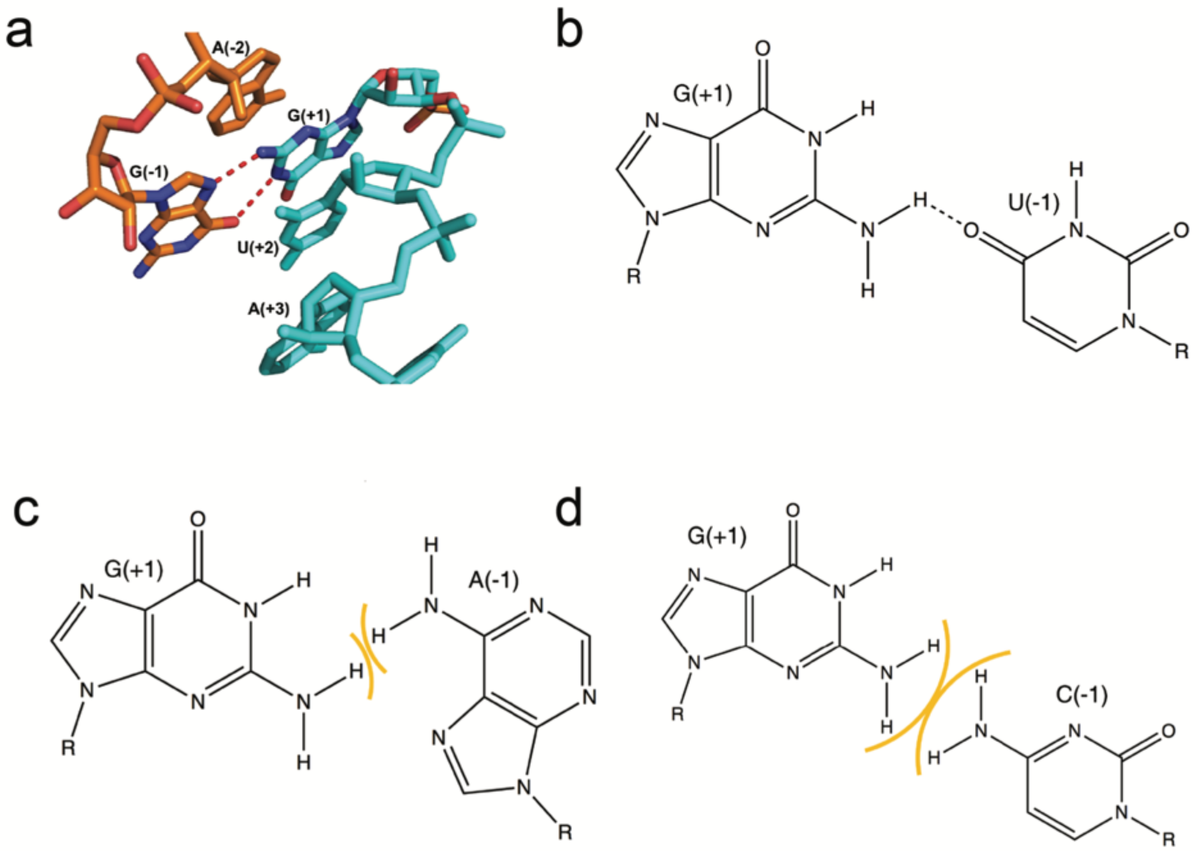

### Supplemental Methods

#### Script Availability

All scripts for the COMPASS read processing pipeline are available on GitHub (https://github.com/k-roy/COMPASS). Scripts used to generate figures for this study available at (https://github.com/k-roy/COMPASS/tree/master/yeast_Prp18).

#### Read 3′ trimming

First, reads were trimmed at the 5′ and 3′ ends for low quality base calls, at 3′ ends for TruSeq adapters, and finally for poly(A) tails by removing T homopolymers at 5′ end of read 1 and A homopolymers at 3′ end of read 2, respectively using cutadapt[1] version 1.18 with the command ***cutadapt --overlap 2 -j 0 -q 20,20 -g "T{100}" -a AGATCGGAAGAGC -A AGATCGGAAGAGC -A "A{100}" -n 2 --trim-n --minimum-length 50 --max-n 4.*** Note that the **--overlap 2** specifies that 2 or more base pairs of matching sequence are needed to trim the adapter. Therefore, a read ending in "TAG" will be trimmed by 2 bases but a read ending in "TTA" will not be trimmed. The option -n 2 instructs cutadapt to look for up to two 3′ adapters, enabling trimming of TruSeq adapters and then poly(A) tails. These trimming operations are important so that the aligners do not need to align terminal stretches of non-genomically encoded bases to the genome. The inclusion of such segments complicates direct comparisons between different aligners, as some aligners may attempt to soft-clip or map terminally with mismatches/indels, while others might search for potential gapped alignments with mismatches/indels. Removing TruSeq adapters, poly(A) tails, and low-quality base calls helps avoid false-positive splice junctions at the level of read mapping and enables more consistent comparisons between aligners.

#### Renumbering read names with integers in chronological order

To simplify comparing each alignment across multiple aligners, read pair names are changed to a simple chronological numbering scheme with the awk command: ***zcat < $IN_DIR$prefix"_R1.fastq.gz" | awk '{print (NR%4 == 1) ? "@" ++i "_R2": $0}' | gzip -c > $OUT_ DIR$prefix"_R1.fastq.gz"*** for read 1 and ***zcat < $IN_DIR$prefix"_R2.fastq.gz" | awk '{print (NR%4 == 1) ? "@" ++i "_R2": $0}' | gzip -c > $OUT_ DIR$prefix"_R2.fastq.gz"*** for read 2 (awk version 20070501).

#### Alignment with BBMap and STAR

Numbered reads were then mapped to the *S. cerevisiae* genome (R64-2-1 version, Saccharomyces Genome Database) with BBMap (version 38.68) (Bushnell, 2014) using parameters **intronlen=20 tipsearch=2000 pairlen=10000,** STAR (version 2.7.0d) (Dobin et al., 2012) using parameters ***--alignEndsType EndToEnd --outSAMattributes NH HI NM MD AS nM jM jI XS*** for the default mode, and with additional options **--scoreGapNoncan 0 --scoreGapGCAG 0 --scoreGapATAC 0** for the non-canonical mode. Note that for a proper comparison of aligned bases at the ends of reads, each aligner must also have an option to disable soft-clipping. This is the default mode for BBMap but this must be activated with STAR using **--alignEndsType EndToEnd**.

Note that the CIGAR fields generated by each aligner must be in 'extended' format (SAM version 1.4) so that a score on each alignment can be directly computed from the CIGAR string. Therefore, each aligner needs to be checked as to whether it outputs matches and mismatches in extended CIGAR format as = and X, respectively (as does BBMap), or as M (as does STAR). In the latter case, the MD tag in the optional fields of the BAM alignment file can be used to generate the extended CIGAR with samfixcigar.jar from Jvarkit (Lindenbaum, 2015). Note that this program will perform substantially faster with coordinate-sorted bam. After converting to extended CIGAR format, the bam files produced from each aligner were sorted by name with samtools version 1.9 (see COMPASS_process_reads_and_align.sh in code repository).

### Comparison of Multiple alignment Programs for Alternative Splice Site discovery (COMPASS)

COMPASS involves four major steps: (1) optimal alignment selection from multiple aligners, (2) alternative splice junction calling, (3) adjustment of ambiguous junctions and (4) junction quality filtering.

#### Optimal alignment selection from multiple aligners

The first step in COMPASS involves comparing alignments for each read across multiple aligners. In this study, we use the BBMap and STAR aligners, although COMPASS will work with any number of alignment programs. The name sorted alignments enable line-by-line comparison of each aligner's bam file, with each read of a read pair appearing on consecutive lines so that the bam files can be read in parallel. The entirety of each bam file therefore does not need to be stored in memory, allowing COMPASS to work with minimal RAM requirements. The chronological numbering allows for determining which aligners did not report a particular read. Each read is first given a score based on the number of bp involved in mismatches, insertions and deletions. Deletions >20 bp are putative introns and are not given a penalty. The scores of each read from a pair are added together to generate the paired-end alignment score. The alignment with the lowest score is selected as the optimal alignment for the COMPASS bam file. The aligner used for a read pair is denoted in the metadata sam field with the program tag PG . Rules for tie-breaking different alignments (i.e. different CIGAR strings) with equal scores are as follows: ungapped alignments are chosen over gapped alignments, and gapped alignments corresponding to annotated junctions are chosen over unannotated junctions. There are several cases where different alignments may receive the same score. For example, a read may map equally well to different loci or there may be equally valid but distinct ways of placing a mismatch or indel in a given region. Furthermore, reads can map to ambiguous junctions where there are identical nucleotides at the end of the upstream exon and end of the intron, or at the beginning of the intron and beginning of the downstream exon. These are cases where the position of the junction could have been shifted upstream or downstream by a constant number of base pairs, respectively, with equal alignment score. To facilitate this step, annotated junctions are pre-processed for the number of ambiguous bp upstream and downstream the junction (relative to + strand). Gapped alignments are considered to map to an annotated junction if they map to an ambiguous adjustment. The outcome of the COMPASS comparison process is designated with: 0 for identical alignments, 1 for a superior alignment score, 2 for equal scores with an ungapped alignment win over gapped alignment, 3 for equal scores with an annotated junction win over unannotated junction, 4 for equal scores but different gapped alignments at the same locus due to ambiguous junction placement (only for unannotated junctions), 5 for equal scores but different alignments at the same locus due to alternate placing of unambiguous junctions, 6 for equal scores but different alignments at the same locus due to alternate placing of mismatches/indels, and 7 for equal scores but different alignments at different loci (i.e. alignments don't overlap). For cases 4-7, the alignment from one aligner is chosen at random; note that the program tag PG still gives each aligner credit for obtaining the best score in this case. The comment tag CO is used to give comma-separated scoring information for each aligner: the aligner name, read 1 CIGAR string, read 2 CIGAR string, the COMPASS alignment score on the read pair, and whether the paired alignment contained annotated splicing junction(s) (ASJ), only unannotated splicing junction(s) (USJ), or only ungapped alignments (UGA). Each aligner entry is separated by semi-colon, and the COMPASS outcome is denoted at the end with a '/' (e.g. **CO:Z:BBMap,100=,80=300N20=,0,USJ;STAR,100=,84=400N2X14=,2,ASJ/1,** where the '/1' denotes a superior alignment is utilized). In this case the program tag would be **PG:Z:BBMap**. If one aligner did not report an alignment for the read, each field after the aligner is denoted by an asterisk. In addition to the optimal alignment bam file, COMPASS outputs separate bam files for ASJ, USJ, and UGA for the optimal alignments to enable viewing alignments of only unannotated splicing junctions in a genome browser. See COMPASS_compare_splice_junctions_from_multiple_aligner_SAM.py in code repository.

#### Alternative splice junction calling

The COMPASS bam file for each sample is then analyzed to create an individual data table for all potential junctions in the sample, including each annotated splice junction as well as each potential alternative splice junction. Each row represents a potential junction, and each column contains sample-specific data (e.g. read support, splicing efficiency, the number of unique gapped alignments [coordinate-CIGAR combos] and corresponding read count supporting each junction) as well as general attributes (e.g. gene name, sequence upstream or downstream of splice sites). For alternative splicing junction filtering, three criteria must be fulfilled: **(i)** The unannotated 5′SS must be on the same strand and situated within 1000 bp of an annotated 5′SS, **(ii)** The alternative intron must at least partially overlap the annotated intron (i.e. the alternative 3′SS must be downstream of the annotated 5′SS, and the alternative 5′SS must be upstream of the annotated 3′SS), and **(iii)** The alternative intron must be ≥20 and ≤2000 bp, consistent with established variation in yeast intron lengths. This can be adjusted depending on the expected intron length distribution for different organisms. All of the split reads supporting each junction are collected, and processed for two elements found to indicate the likelihood of the junction corresponding to a *bona fide* splicing event – the longest segments on either side of the putative junction mapped without a mismatch, and the fraction of the mapped reads harboring a junction-proximal mismatch (where junction proximal is defined as ≤10 bp from the junction).

#### Adjustment of ambiguous junctions

Ambiguous junctions are identified by the presence of identical nucleotides at the end of the upstream exon and end of the intron, or at the beginning of the intron and beginning of the downstream intron. These are cases where the position of the junction could have been shifted upstream or downstream, respectively, with equal alignment score. In this step the 5′SS consensus sequence (GUAUGU in the case of budding yeast) is used to resolve junction ambiguity. The junctions are adjusted so that the 5′SS with the closest match to the consensus is utilized. The coverage at the 5′SS and 3′SS in the UGA files correspond to the unspliced read signal and is used to calculate the annotated splicing efficiency (annSE) and alternative splicing efficiency (altSE). The fraction of annotated splicing (FAnS) is calculated as the ratio of alternative to annotated splice junction reads. See COMPASS_integrate_splice_junction_profiles_from_multiple_samples.py.

#### Junction quality filtering

The next step in COMPASS is to integrate the individual sample tables into a combined data table. Statistics across all samples are calculated for each putative junction, including the longest upstream and downstream overhangs without a mismatch (i) the fraction of gapped reads without a mismatch within 10 bp of the junction, and the longest perfect overhang among all supporting reads upstream (ii) and downstream (iii) of the junction. To establish appropriate metrics for junction filtering, these metrics were plotted by total read support with annotated and unannotated junctions colored separately (as shown in Figs S3 and S4). In this study, we determined appropriate cutoffs for each junction to be (i) 50% of supporting reads without a mismatch within 10 bp of the junction, (ii) longest perfect overhang ≥10 bp for upstream the 5′SS, (iii) longest perfect overhang ≥40 bp for downstream the 3′SS. These thresholds can be adjusted by the user. See COMPASS_analyze_splice_junction_profiles_for_individual_samples.py.
